# A spatiotemporal transcriptomic atlas of the mouse placenta reveals glycogen cell-mediated metabolic support essential for fetal viability

**DOI:** 10.1101/2024.05.28.596242

**Authors:** Yuting Fu, Xiaoqi Zeng, Yifang Liu, Shikai Jia, Yujia Jiang, Jia Ping Tan, Yue Yuan, Tianchang Xia, Yun Mei, Shan Wen, Xiaojing Liu, Yue You, Weike Pei, Chengshuo Yang, Sida Shao, Junhua Shen, Liangshan Mu, Xiaoxue Ma, Matthew Paul McCormack, Saifeng Cheng, Luyi Tian, Longqi Liu, Xiaoyu Wei, Xiaodong Liu

## Abstract

Proper placentation is fundamental to the growth and viability of the maturing embryo. However, a thorough grasp of the spatial organization of cell types, their interactions, and gene expression patterns along the maternal-fetal function during placental development remains incomplete. Here, we utilized Stereo-seq to construct the spatiotemporal transcriptomic atlas of the mouse placenta (STAMP) spanning embryonic (E) days 9.5 to 18.5 at single cell resolution (https://db.cngb.org/stomics/stamp/). This resource delineates spatially resolved cellular dynamics and gene expression patterns across placental and maternal compartments. We identified distinct glycogen trophoblast cell (GC) subclusters, mapped their developmental trajectories, and uncovered transcriptional transitions accompanying their migration from the junctional zone to the maternal decidua from E12.5 onward. In a defective placentation model with perinatal lethality, GCs abnormally persisted at E18.5 with excessive glycogen and reduced degradation metabolites in placenta and fetal liver, indicating impaired glycogen breakdown. Maternal glucose supplementation restored glucose levels and rescued fetal survival, underscoring GC-mediated metabolic support as critical for viability. Together, this study provides a comprehensive spatiotemporal placental atlas and demonstrates that GC-mediated glycogen metabolism is essential for sustaining fetal viability.

## Introduction

The placenta, a sophisticated and unique organ, plays an indispensable role during mammalian pregnancy. It not only serves as a physical barrier between maternal blood and the fetus, but also performs crucial functions such as delivering oxygen and nutrients to the fetus and removing waste^1,2,3^. A dysfunctional placenta is often linked to various human pregnancy-related disorders, including miscarriage, preeclampsia, and fetal intrauterine growth restriction^4,5,6^.

Morphological and functional similarities between human and mouse placenta make the mouse a key model for studying the impact of placenta development on pregnancy and the mechanisms underlying placental dysfunction^7,8^. The mature mouse placenta comprises the maternal decidua basalis and the fetal-derived labyrinth and junctional zone^9,10^. The labyrinth mediates nutrient and gas exchange, and its disruption often leads to developmental failure and growth restriction, highlighting its essential role in nutrient transfer from mid-gestation onward^11,12^. By contrast, the junctional zone is mainly endocrine and consists of spongiotrophoblast (SpT) and glycogen trophoblast cells (GC)^13^. GCs are named for their storage of glucose as glycogen, although the physiological significance of this storage remains unclear^14^. Beginning around E12.5, immature GCs gradually mature, accumulating glycogen and acquiring a vacuolated morphology, with GC numbers increasing nearly 300-fold between E12.5 and E16.5 before declining by approximately 50% by E18.5^15^. In humans, aberrant placental glycogen storage is associated with maternal diabetes and pre-eclampsia^16,17,18,19^, and over 40 targeted mouse mutations demonstrate that defects in GCs compromise fetal growth^14^, suggesting placental glycogen may provide readily mobilized glucose during periods of high fetal demand^15,20^. However, direct functional evidence is still lacking. Importantly, up to 68% of knockout mouse lines that are lethal at or after mid-gestation exhibit placental abnormalities, and early embryonic deaths between E9.5 and E14.5 are almost invariably associated with severe placental malformations^11^. These defects correlate strongly with abnormal brain, heart, and vascular development^21^, highlighting the critical need to elucidate the cellular and molecular mechanisms by which placental dysfunction compromises normal embryonic growth and viability.

Previous studies have uncovered cell types comprising trophoblast derived lineages of mouse placentas at various developmental stages (from E7.5 to E14.5) using single cell/nucleus RNA-sequencing^22,23^. Yet, our understanding of placental spatial organization, molecular dynamics, and functional maturation, and of how their disruption affects embryonic development, remains limited. Here, we reconstructed a spatiotemporal atlas of mouse placentation across E9.5 to E18.5 using Stereo-seq, the data is accessible via interactive data portal, named spatiotemporal transcriptomic atlas of mouse placenta (STAMP) (https://db.cngb.org/stomics/stamp/). This resource advances insight into placental ontogeny and cell-cell interactions at the maternal-fetal interface and establishes a reference for evaluating genetic or pathological perturbations. Mining this atlas revealed that glycogen trophoblast cells progressively migrate from the junctional zone to the maternal decidua while undergoing transcriptional state changes and forming previously unrecognized subclusters with distinct molecular profiles. Application to a defective placentation model with perinatal lethality demonstrated that disruption of glycogen metabolism compromises fetal survival. These findings highlight placental glycogen metabolism as a fundamental requirement for sustaining fetal viability.

### A spatiotemporal single-cell transcriptomic atlas of mouse placentation

To gain a comprehensive understanding of cell fate transitions and specifications at spatial and temporal resolution during placentation, we first profiled 9 mouse placentas spanning developmental stages from E9.5 to E18.5 using Stereo-seq and snRNA-seq (Figure 1A, S1A, S1B and S1C). After rigorous quality control procedures (see Methods), we identified 35 distinct cell clusters, broadly categorized into trophoblast cells (14 subclusters), stromal cells (4 subclusters), immune cells (7 subclusters), endothelial cells (4 subclusters), and other cell types from both fetal and maternal tissues (Figure 1B, S1D, and Supplementary Table 1)^22,23,24,25,26^. We validated and confirmed these cell annotations by cross-referencing them with existing placental datasets from prior studies (Figure S1E)^22,23^.

**Figure 1.**
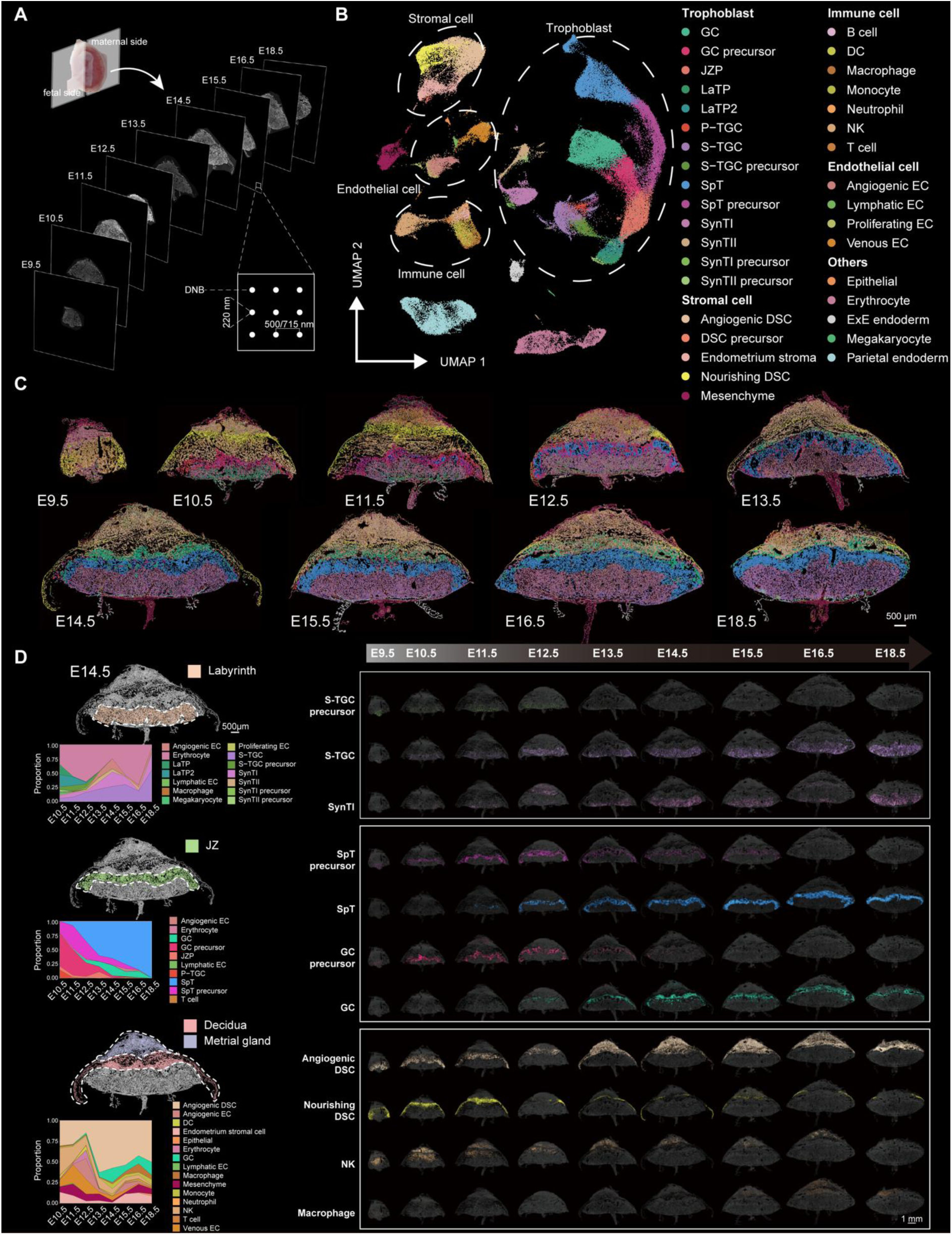
**A spatiotemporal single-cell transcriptomic atlas of mouse placentation (E9.5–E18.5)** (A) Schematic illustration of the placenta samples collected for spatial transcriptomics profiling. (B) Uniform manifold approximation and projection (UMAP) of snRNA-seq clustering showing 35 cell types. GC, glycogen cell; JZP, junctional zone progenitor; LaTP, labyrinth trophoblast progenitor; P-TGC, parietal trophoblast giant cell; S-TGC, sinusoid trophoblast giant cell; SpT, spongiotrophoblast cell; SynT, syncytiotrophoblast cell; DSC, decidual stromal cell; EC, endothelial cell; DC, dendritic cell; NK, natural killer cell. (C) Spatial visualization of all cell distributions within placentas at different developmental stages from E9.5 to E18.5 using Stereo-seq data. Cells are colored by their annotations. Scale bars, 500 μm. (D) Cell type distribution and quantification in placenta sections from E9.5 to E18.5. Anatomical regions (left) were identified as Labyrinth zone, Junctional zone (JZ), Maternal decidua, and Myometrium. Regions are colored based on anatomical region annotations. Cells are colored based on cell type annotations. Stacked area plots showing cell type proportions in the corresponding regions. Scale bars, 500 μm.

Next, we employed DNA staining combined with the watershed algorithm^27^ to achieve single-cell resolution, enabling the capture of transcripts within clearly defined nuclear and cytoplasmic boundaries. Through spatially constrained clustering (SCC), we delineated distinct clusters corresponding to key anatomical regions, including the labyrinth, junctional zone (JZ), maternal decidua, and metrial gland (Figure S2A). Using snRNA-seq data as a reference, we annotated the Stereo-seq dataset with the TACCO (v0.3.0) framework^28^, assigning the most probable cell type to each individual cell (see Methods for details). This strategy allowed us to comprehensively map the spatial and temporal transitions of placental cell types across developmental stages from E9.5 to E18.5. The full dataset is publicly available on our interactive platform, the Spatiotemporal Transcriptomic Atlas of Mouse Placenta (https://db.cngb.org/stomics/stamp/) (Figure 1C, 1D, S2B, S2C and S2D).

Firstly, our study delineates distinct spatial distributions and dynamic fate transitions among placental cell types across key anatomical regions, illuminating their region-specific functions and overall significance during embryonic development. In the labyrinth region, we observed expansion through branching morphogenesis that creates intricate vascular spaces essential for maternal-fetal blood exchange^29^. Notably, the proportion of trophoblast progenitor cells in the labyrinth decreases steadily from E9.5, while terminally differentiated trophoblast cells surge after E12.5, marking critical milestones in placental maturation (Figure 1C and 1D). Simultaneously, spongiotrophoblast (SpT) and glycogen trophoblast cells (GCs) begin to mature from E12.5 onward.

Remarkably, we detected a spatial transition of GCs from the junctional zone (JZ) to the maternal decidua. GCs first appear in the JZ at E12.5, peak at E14.5, and then decline, while their emergence in the decidua begins at E13.5 and increases steadily until E18.5 (Figure 1C and 1D). This pattern corroborates previous immunostaining studies using protocadherin 12 (PCDH12) as a GC marker^15,30^. In contrast, other cell types, including endothelial cells, immune cells, and stromal cells are predominantly localized within the maternal decidua (Figure 1C and 1D).

Leveraging our spatial transcriptomic dataset, we further identified multinucleated syncytiotrophoblast cells and trophoblast giant cells (TGCs) across multiple developmental stages *in situ* (Figure 1C and 1D), a feat previously unattainable with conventional single-cell or single-nucleus RNA-seq methods^22,23^. Additionally, we captured the spatial dynamics of two distinct decidual stromal cell (DSC) populations: angiogenic DSCs and nourishing DSCs, each exhibiting unique molecular signatures and functions^24^. From E9.5, angiogenic DSCs migrate toward the upper decidua, near the myometrium, to promote angiogenesis, stabilizing in number after E14.5. Conversely, nourishing DSCs, which are involved in energy homeostasis and hormone metabolism^24,31^, gradually decline and shift toward the fetal side of the placenta (Figure 1C and 1D).

In summary, by integrating spatial transcriptomics and snRNA-seq datasets, we decoded spatially resolved transcriptomes at single-cell resolution and determined cell type composition during mouse placentation. Crucially, the dynamic spatial changes of cell types over time illuminated key developmental milestones during placentation.

### Molecular and spatial characterization of glycogen trophoblast cell subtypes

We next examined the spatiotemporal development of trophoblast cells, the primary cell types of the placenta. Reclustering the snRNA-seq data revealed two distinct glycogen cell (GC) subclusters (Figure 2A and Supplementary Table 2). Mapping their spatial distributions using Stereo-seq data showed that one subcluster (GC-1) was predominantly in the junctional zone (JZ), while the other (GC-2) was confined to the maternal decidua (Figure 2B).

**Figure 2.**
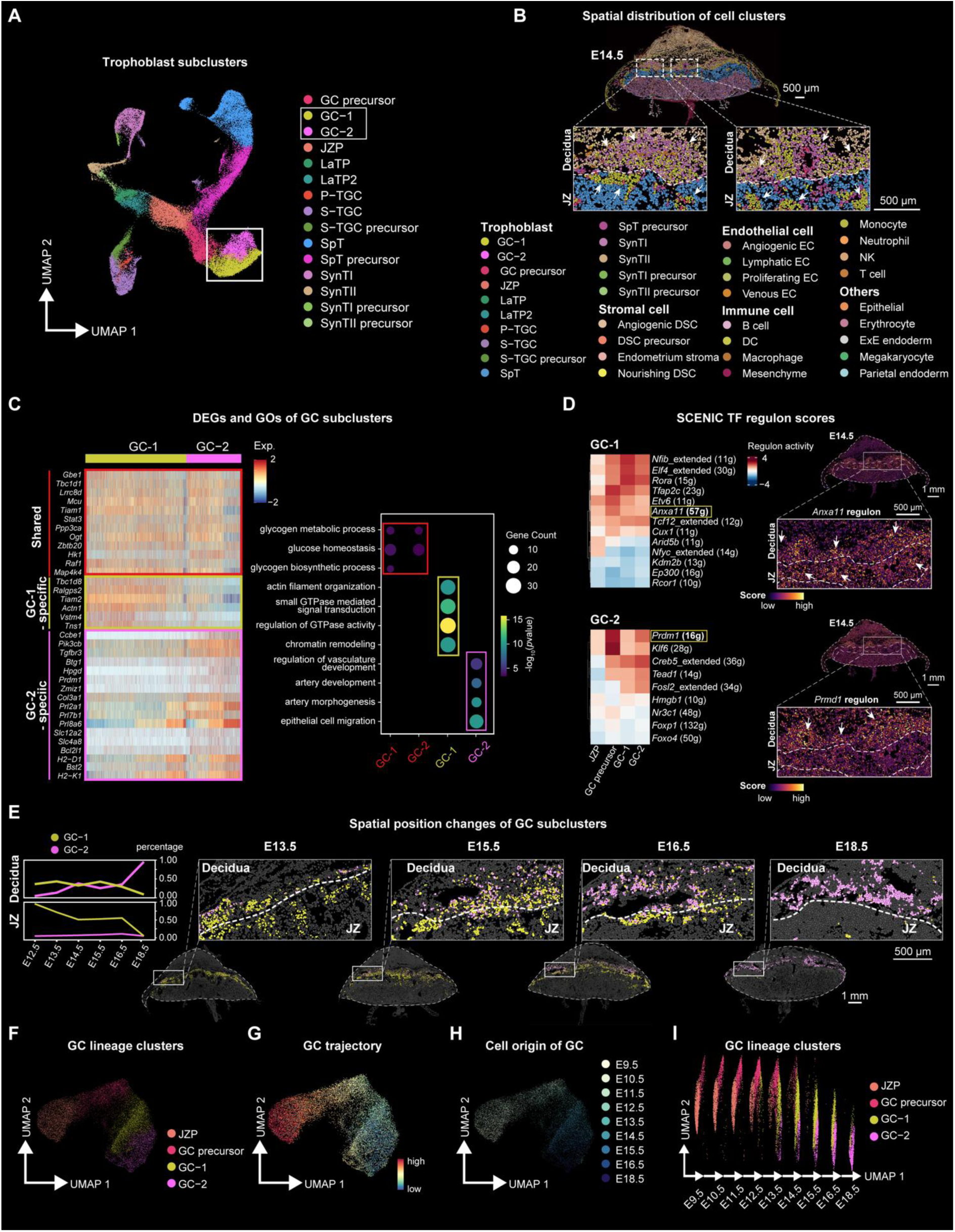
**Molecular, spatial, and developmental characterization of GC subtypes** (A) UMAP visualization showing the reclustering of trophoblast cells. (B) Spatial visualization of all cell distributions in the E14.5 placenta section. Two magnified fields of view highlight the distinct spatial distribution of the two GC subclusters. Scale bars, 500 μm. (C) Heatmap (left) showing the DEGs of GC subclusters, with significantly enriched GOs shown by bubble plot (right). Shared genes and GOs are outlined in red, GC-1-specific genes and GOs are outlined in yellow, and GC-2-specific genes and GOs are outlined in pink. (D) Heatmap (left) showing the SCENIC transcription factor (TF) regulon scores in GC-1 and GC-2 based on regulon activity. Spatial visualization of selected regulons, *Anxa11* and *Prmd1*, are shown respectively in the E14.5 placenta section (right). (E) Line chart (left) showing the percentage of the two GC subclusters in the maternal decidua and JZ region from E12.5 to E18.5. The spatial visualization of the two GC subclusters at E13.5, E15.5, E16.5 and E18.5 is shown on the right. Cells are colored by their annotations. Scale bars, 500 μm. (F) UMAP visualization showing cell types across GC differentiation, including junctional zone progenitor (JZP), GC precursor, and two GC subclusters. Cells are colored by cell type annotation. (G) Pseudotime trajectory of cell types across GC differentiation, analyzed using Monocle3 and CytoTRACE. Cells are colored by pseudotime. (H) UMAP visualization showing the origin of GC based on the time point. (I) UMAP visualization of cell types across GC differentiation. Cells are colored by cell type annotation.

To dissect the molecular differences between these GC subclusters, we performed differential gene expression analysis (Figure 2C and Figure S3A; Supplementary Table 3). *Aldh1a3*, a known marker of glycogen cells and their precursors^32^, was highly expressed in GC-1, whereas *Prl7b1*, overlapping with *Pcdh12* expression in the maternal decidua^33^, was more abundant in GC-2. These patterns were corroborated by their spatial distributions (Figure S3B), and were further confirmed by RIBOseq analysis^34^, which provided spatially resolved measurements showing predominant *Aldh1a3* and *Prl7b1* protein synthesis in the E14.5 and E18.5 placental sections (Figure S3C). Gene ontology analysis further revealed that GC-1 was enriched in processes such as chromatin remodeling and GTPase-mediated signal transduction, while GC-2 was associated with epithelial cell migration and vascular development (Figure 2C; Supplementary Table 4). Notably, genes shared between GC-1 and GC-2 were enriched in glycogen metabolism pathways (Figure 2C; Supplementary Table 4).

To explore the transcriptional dynamics underlying GC fate, we conducted SCENIC analysis to assess transcription factor (TF) expression and motif enrichment. This approach identified key regulons with differential activity between the subclusters, including established TFs (*Tfap2c*, *Tcf12*, *Prdm1*, *Fosl2*)^35,36,37,38^ as well as novel ones (*Nfib*, *Elf4*, *Etv6*, *Anxa11* in GC-1; *Creb5* in GC-2). Spatial transcriptomics validated these regulons in E14.5 placenta sections (Figure 2D and S3D). Furthermore, using the CellChat algorithm^39^, we uncovered distinct cellular communication networks: GC-2 showed signaling interactions (e.g., *Sema3e*-*Plxnd1*) indicative of an angiogenic role^40^, as well as pathways involving *Lgals9* and receptors such as *Ptprc*, *Ighm*, *Havcr2*, and *Cd44*, suggesting immunomodulatory functions. Additionally, *Ccl27a*, which is involved in immune cell recruitment, was predominantly expressed in GC-2, while both subclusters secreted growth factors like *Igf2* (Figure S4E and S4F).

Temporal trajectory analysis of the GC lineage revealed that GC-1 emerges early in the JZ, peaks briefly, and then declines, whereas GC-2 appears after E13.5, becomes restricted to the maternal decidua, and increases steadily until E18.5 (Figure 2E and S4G). Overall GC numbers rise from E12.5 to E16.5 before markedly decreasing by E18.5 (Figure 2E and S4G), consistent with reports of extensive GC lysis during the perinatal period^15,41^. Trajectory reconstruction using Monocle3^42^ and CytoTRACE^43^ further delineated a developmental pathway following a JZ progenitor – GC precursor – GC-1 – GC-2 sequence (Figure 2F, 2G, 2H, 2I, and S4A).

In summary, our findings reveal previously uncharacterized, transitioning GC subtypes with distinct spatial distributions, molecular profiles, and regulatory networks, and uncovered transcriptional transitions accompanying their migration from the junctional zone to the maternal decidua from E12.5 onward.

### Trajectory analysis reveals regulators of GC lineage linked to embryonic lethality

Embryonic lethality in mice is often linked to impaired placental function^11^. To investigate this connection, we compiled a comprehensive list of 151 mouse mutant lines associated with embryonic death and placental development defects. This list includes 103 knockout lines from the Deciphering the Mechanisms of Developmental Disorders (DMDD) program (https://dmdd.org.uk) and an additional 48 mutant lines from other studies^11^. We categorized these genes by the timing of lethality and reviewed the literature for evidence of placental abnormalities (Table 1).

**Table 1.** All lethal mouse knockout lines according to previous reports. All genes are categorized according to the time of lethality. They are roughly divided into postnatal lethality and lethality at various stages of embryonic development. The table also provides information on whether placental phenotype defects have been reported, along with the corresponding references. Lethal, no homozygotes are recovered. Subviable, homozygotes recovered but at less or equal to 13% (≤13%). Viable, homozygotes recovered more than 13% (>13%).

To further elucidate placental development, we performed trajectory analysis of trophoblast cells using Monocle3 and CytoTRACE^42,43^, which revealed five principal trophoblast lineage progression trajectories, including the GC lineage (Figure S4A and S4B). Along these trajectories, we identified genes with temporally regulated expression patterns along pseudotime that likely drive key stages of placental development (Figure S4C)^44^.

We identified *Ano6*, a member of the Anoctamin chloride channel family, as a crucial factor in GC lineage development and placental function (Figure 3A, 3B, S5A, and S5B), with previous studies showing that *Ano6*-deficient mice exhibit impaired placental development and perinatal lethality^45^. RT-PCR analysis indicates that *Ano6* is uniquely expressed and upregulated in the placenta (Figure S6A). SnRNA sequencing data further reveal that *Ano6* is specifically highly expressed in GCs at various developmental stages (Figure S6B and S6C), a finding that is also corroborated by RIBOseq experiments, which provided spatially resolved measurements showing predominant *Ano6* protein synthesis in the junctional zone (JZ) at embryonic day 13.5 (E13.5) (Figure 3C).

**Figure 3.**
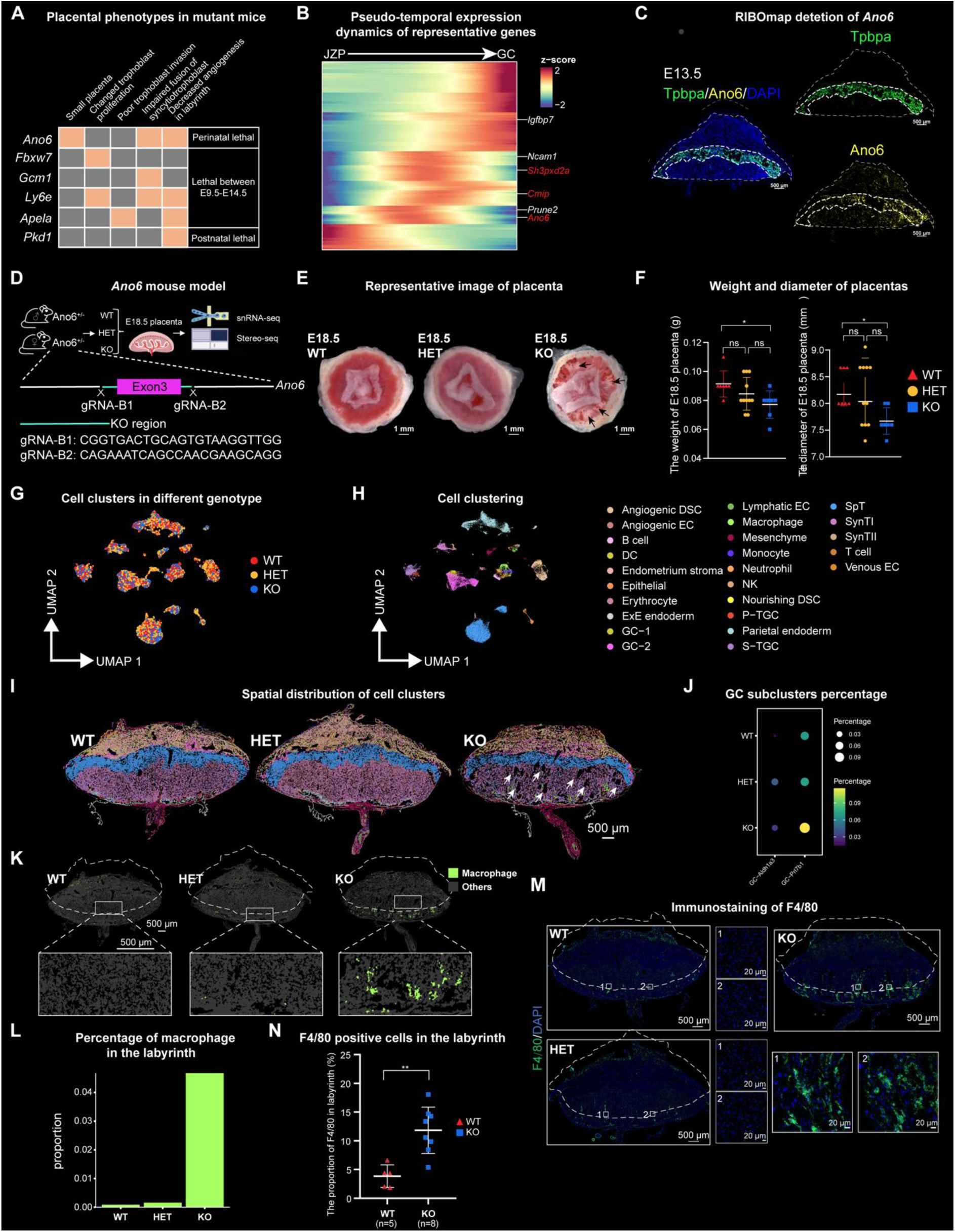
***Ano6* deficiency disrupts placental development and increases GC abundance** (A) Representation of the curated list of embryonic lethal mouse mutant genes (orange denotes abnormality detected). The selected mutant genes are categorized by lethality stage and marked with corresponding placental phenotypes. (B) Pseudo-temporal expression dynamics of specific representative genes along the GC developmental trajectory. Genes previously identified to be associated with lineage development are labeled in black, while potential regulators whose loss leads to embryonic lethality are highlighted in red. (C) Detection of *Ano6* and *Tpbpa* protein synthesis by RIBOseq in E13.5 placenta sections. Dotted lines encircle the same region. Scale bars, 500 μm. (D) Schematic illustration of the experimental design. (E) Representative images of placentas from WT, HET, and KO mice at embryonic day 18.5 (E18.5). Black arrows point to the phenotypic defects in the KO placenta. *Ano6* KO placentas are smaller than WT and HET, and exhibit prominent white plaques on the fetal side, indicative of vascular structural defects. (F) Quantification of the weight and diameter of the WT, HET, and KO placenta. WT (*n*=7), HET (*n*=11), KO (*n*=7). One-way analysis of variance (ANOVA). \**p* < 0.05. All data represent means ± SEM. (G) UMAP representation of all cell types in E18.5 WT, HET, and KO placentas. (H) UMAP visualization of snRNA-seq clustering showing the 25 cell types in E18.5 WT, HET, and KO placenta samples. (I) Spatial visualization of all cell distributions in Stereo-seq data. Cells are colored by their annotations. Scale bars, 500 μm. (J) Bubble plot showing the percentage of GC subclusters in E18.5 WT, HET, and KO placentas. (K) Spatial visualization of macrophages in E18.5 WT, HET, and KO placentas. Cells are colored by their annotations. Scale bars, 500 μm. (L) Bar plot showing the percentage of macrophages in the labyrinth of E18.5 WT, HET, and KO placentas. (M) F4/80 immunofluorescence staining of E18.5 WT, HET, and KO placentas. Cross sections of the entire placentas and enlarged views are shown, respectively. Scale bars, 500 μm (overall shape) and 20 μm (high-magnification view). (N) The proportion of F4/80-positive cells in the labyrinth is quantified in WT and KO placentas. WT (n=5), HET (n=5), and KO (n=8). One-way analysis of variance (ANOVA). \*\**p* < 0.01, \*\*\**p* < 0.001. All data represent means ± SEM.

### Spatial transcriptomic profiling of placental defects in an embryonic-lethal mutant

To uncover the mechanisms behind placental defects linked to embryonic lethality, we generated *Ano6* knockout (KO) mice by deleting exon 3 of the *Ano6*-208 transcript, resulting in a frame-shift mutation (Figure 3D, S6D and S6E). Due to the low postnatal survival rate of homozygous KO mice (3.03%) (Table 2), we bred heterozygous pairs to obtain KO offspring. Compared with wildtype (WT) and heterozygous (HET), KO placentas were smaller and lighter, and displayed prominent large white plaques on the fetal side, suggesting defects in vascular formation (Figure 3E and 3F), consistent with the previous report^45^. The phenotypic differences among WT, HET and KO placentas became apparent at embryonic day 18.5 (E18.5) (Figure S6F and S6G). Specifically, KO placentas exhibited labyrinth defects characterized by enlarged cavities, reduced area and cellularity, and impaired vascular structure (Figure S6H-M), consistent with previously reported phenotypes^45^.

**Table 2.** Genotype distribution of Ano6 knockout mice.

To probe molecular differences *in situ*, we combined Stereo-seq with snRNA-seq analysis of WT, HET, and KO samples at E18.5 (Figure S7A and S7B). This integrated approach identified various cell clusters, including trophoblasts, immune cells, stromal cells, and endothelial cells (Figure 3G, 3H, 3I, and S7C). Notably, KO placentas at E18.5 contained markedly increased numbers of glycogen trophoblast cells (Figure 3J and S7D), indicating potential abnormalities in their development or function. While total macrophage numbers were similar across genotypes (Figure S7D), macrophages were enriched in the labyrinth layer of KO placentas (Figure 3K, 3L, and S7E), as confirmed by F4/80 immunostaining (Figure 3M and 3N).

In summary, in addition to phenotypes consistent with previous reports, our knockout placentas at E18.5 showed increased glycogen trophoblast cells, and abnormal macrophage accumulation in the labyrinth, suggesting that impaired glycogen metabolism and aberrant immune responses may underlie placental dysfunction and perinatal lethality.

### GC persistence and impaired glycogen catabolism in *Ano6*-null placentas

The increased number of glycogen cells in KO placentas was confirmed by Periodic acid-Schiff (PAS) and hematoxylin and eosin (H&E) staining (Figure 4A, 4B, S8A, and S8B). Under normal physiological conditions, glycogen trophoblast cells (GCs) expand nearly 300-fold by E16.5 but decline markedly by E18.5 as they undergo lysis/apoptosis and form glycogen-filled lacunae near vascular sinuses, a process thought to supply energy at term or influence parturition^30^. Since GCs serve as glycogen reservoirs, we next used transmission electron microscopy (TEM) to visualize glycogen granules (Figure 4C). Quantitative analysis showed that KO GCs contained significantly more glycogen granules than those in WT and HET placentas (Figure 4D and 4E), indicating impaired glycogen metabolism. Consistently, total placental glycogen content was significantly elevated in KO placentas compared with WT and HET controls (Figure 4F). Adjustment for placental weight yielded similar results, excluding placental size as the explanation for this increase (Figure 4G). Time-course analysis of total placental glycogen from E12.5 to E18.5, using both quantitative measurement and PAS staining of placental sections, further demonstrated that this phenotype emerged only at E18.5, when KO placentas exhibited significantly higher glycogen content than WT and HET placentas (Figure S8C, S8D and S8E).

**Figure 4.**
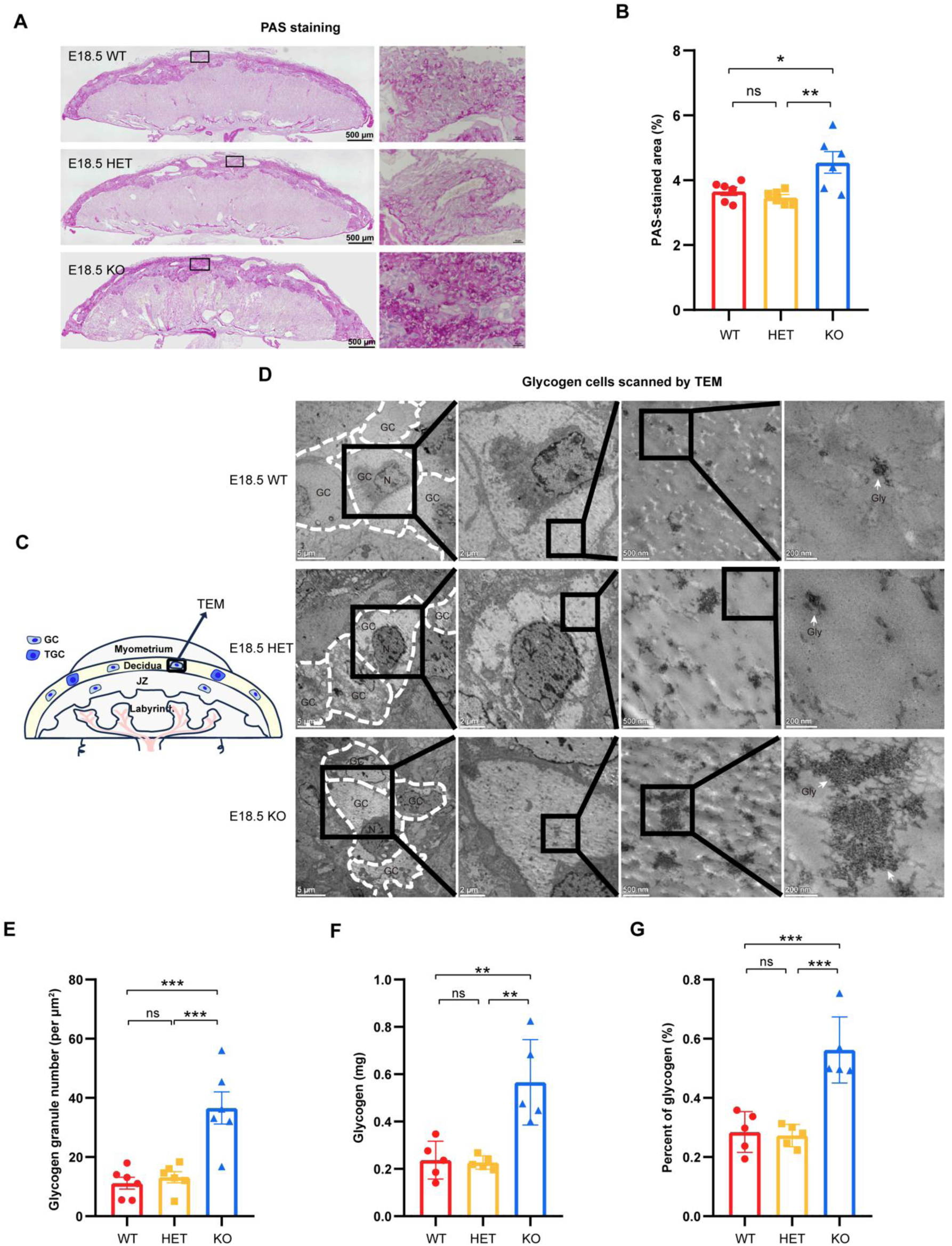
**GC persistence and excessive glycogen accumulation in *Ano6*-null placentas** (A) Overall shape and high-magnification images of the PAS-stained WT, HET and KO placentas. Scale bars, 500 μm (overall shape) and 10 μm (high-magnification view). Quantification was performed on GCs located in both the junctional zone (JZ) and decidua. (B) Percentage of PAS-stained positive area relative to the total tissue area analyzed using ImageJ. WT (*n*=6), HET (*n*=6), KO (*n*=6). One-way analysis of variance (ANOVA). \**p* < 0.05, \*\**p* < 0.01. All data represent means ± SEM. (C) Schematic illustration of glycogen cells scanned by transmission electron microscope (TEM). (D) Electron micrographs of WT, HET, and KO placentas at E18.5. The outline of glycogen cells is circled with white dashed lines. GC, glycogen cells; N, nucleus; Gly, glycogen granules. Scale bars, 500 μm (column 1), 200 μm (column 2), 500 nm (column 3), and 200 nm (column 4). (E) Glycogen granule density in WT, HET, and KO placentas at E18.5. Glycogen granule number per μm^2^ was calculated. WT (*n*=6), HET (*n*=6), KO (*n*=6). One-way analysis of variance (ANOVA). \*\*\**p* < 0.001. All data represent means ± SEM.(F) Total placental glycogen content (mg) in WT, HET, and KO placentas at E18.5. WT (*n*=6), HET (*n*=6), KO (*n*=6). One-way analysis of variance (ANOVA). \**p* < 0.05. All data represent means ± SEM. (G) Placental glycogen expressed as a percentage of placental weight in WT, HET, and KO placentas. WT (*n*=6), HET (*n*=6), KO (*n*=6). One-way analysis of variance (ANOVA). \*\**p* < 0.01. All data represent means ± SEM.

In summary, at E18.5, KO placentas showed increased GCs with excessive undegraded glycogen granules, indicating disrupted glycogen catabolism.

### Defective GC glycogen degradation reduces fetal energy supply and compromises survival

To confirm defective glycogen breakdown in KO placentas, we measured key intermediates of glycogenolysis in both placental and fetal liver tissues by LC-MS analysis, including glucose-1-phosphate (G1P), glucose-6-phosphate (G6P) and glucose (Figure 5A and 5B). KO placentas showed markedly reduced levels of these metabolites (Figure 5C), and a similar reduction was also observed in fetal liver (Figure 5D), supporting the idea that impaired glycogen degradation in placental GCs compromises fetal energy supply and contributes to perinatal lethality. During late gestation, when energy demand is high^15,20^, the sharp decrease in G6P and glucose suggests insufficient metabolic fuel to sustain placental and fetal development^14,30,46^. To test this hypothesis, pregnant dams received daily oral glucose gavage from E13.5 to E18.5 (Figure 5E). Glucose supplementation significantly improved KO fetal survival (from 3.03% to 10.8%) and restored key metabolites, particularly energetically favorable metabolites G6P and glucose, in both placenta and fetal liver (Figure 5F, 5G, 5H, and Table 3).

**Figure 5.**
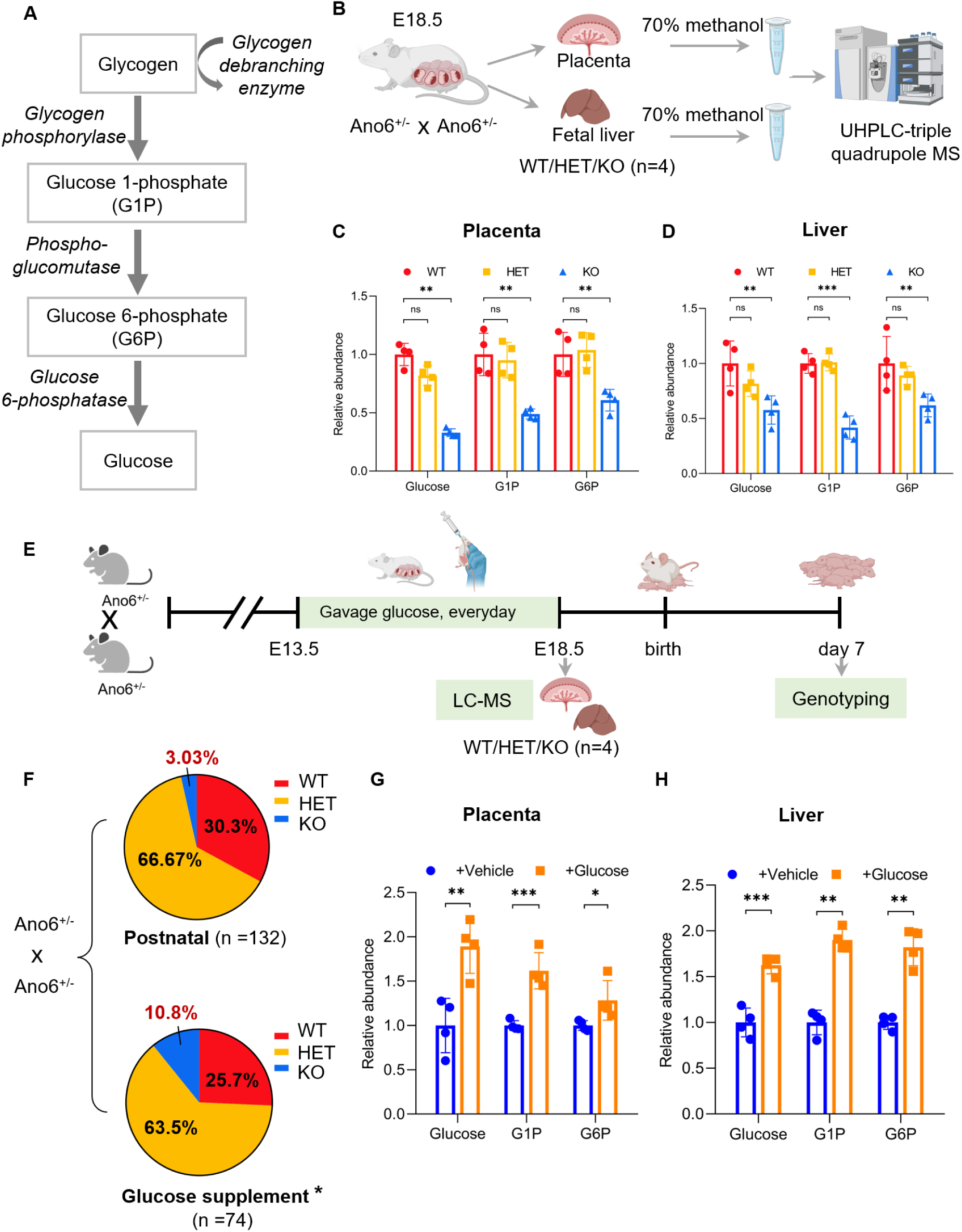
**Defective GC glycogen degradation reduces fetal energy supply and survival** (A) Schematic of glycogenolysis pathway. (B) Schematic workflow for metabolites analysis in placenta and fetal liver by LC-MS. (C and D) Relative abundance of Glucose, G1P and G6P in placental (C) and fetal liver tissues (D) measured by targeted metabolomics. Data are presented as mean ± SEM from four independent experiments. Statistical significance was assessed using One-way analysis of variance (ANOVA). \*\**p* < 0.01, \*\*\**p* < 0.001. (E) Schematic of maternal glucose supplementation strategy. (F) Significant increase survival of *Ano6*^-/-^ mice after maternal glucose supplementation. * *p* < 0.05, Fisher’s exact test. (G and H) Relative abundance of Glucose, G1P and G6P in placental (G) and fetal liver tissues (H) after glucose supplementation measured by targeted metabolomics. Data are presented as mean ± SEM from four independent experiments. Statistical significance was assessed using One-way analysis of variance (ANOVA). \**p* < 0.05, \*\**p* < 0.01, \*\*\**p* < 0.001.

**Table 3.** Genotype distribution of Ano6 knockout mice after glucose supplementation with pregnant mice.

To investigate the cause of abnormal glycogen metabolism in KO placentas, we examined GC differentiation and the expression of glycogenolytic enzymes^13^. GC marker genes, *Pchd12*, *Aldh1a3* and *Gjb3*, showed comparable expression among WT, HET, and KO placentas from E13.5-E18.5 (Figure S9A), indicating intact GC differentiation. Likewise, key glycogenolytic enzymes were expressed at similar levels across genotypes (Figure S9B and S9C). These findings suggest that the metabolic defects in KO placentas are not due to impaired GC differentiation or transcriptional downregulation of glycogenolytic enzymes, but likely reflect a functional impairment in glycogen utilization.

Together, these results demonstrate that loss of *Ano6* blocks glycogen breakdown in placental GCs, resulting in metabolic insufficiency that underlies placental dysfunction and perinatal lethality, while maternal glucose supplementation can partially rescue this defect by restoring energetically favorable metabolites.

### Secondary macrophage accumulation in the labyrinth of *Ano6*-null placentas

In addition to the GCs-related defects identified above, previous studies have reported that *Ano6* deficiency also causes defective trophoblast syncytialization in the SynT-2 layer, leading to maternofetal exchange deficiency, labyrinth malformation, and perinatal lethality^45^. These findings suggest that multiple pathological processes may converge to cause placental insufficiency in *Ano6*-null mice.

To further leverage the strength of spatial transcriptomics in capturing *in situ* microenvironmental changes that are often missed by dissociative single-cell approaches, we next explored cell type distributions within the labyrinth. Interestingly, our spatial data revealed a pronounced accumulation of macrophages specifically in the labyrinth region of KO placentas, which aligns with the known labyrinthine structural defects and highlights the added value of our spatial atlas in detecting region-specific cellular alterations.

Differential gene expression analysis of the labyrinth layer identified 521 genes upregulated and 253 genes downregulated in KO placentas (adjusted *p*<0.05, |log2FC|>1) (Figure S10A; Supplementary Table 5). Upregulated genes were enriched for innate immune response and inflammatory pathways, including Toll-like receptors (*Tlr7*, *Tlr13*)^47^, chemokine ligands (*Ccl4*, *Ccl6*), and immunomodulatory molecules (*Sirpa*, *Ccr5*, *Spp1*, *Ptafr*, *Trem2*) (Figure S10A, S10B and Figure S11)^48,49,50,51,52,53,54^, whereas several solute carrier transporters (*Slc7a2*, *Slc22a4*, *Slc28a3*) were downregulated (Figure S10A, 10C, and Figure S12). Gene ontology analysis confirmed enrichment of terms related to TNF production, leukocyte migration, and tissue remodeling among the upregulated genes (Figure S10D and Figure S11).

Focusing on macrophages, we found 140 genes significantly upregulated in labyrinthine macrophages compared with maternal-side macrophages (adjusted *p*<0.05, |log2FC|>1) (Figure 6A; Supplementary Table 6), including tissue remodeling genes (*Spp1*, *Mmp12*, *Ctss*, *Gpnmb*) (Figure 6B and 6C)^55,56,57,58,59^. Immunostaining showed that F4/80⁺ macrophages co-expressed TGFβ (Figure 6D and S13A), indicating an activated state potentially involved in local immune regulation and tissue repair^60^. Their spatial distribution is closely associated with areas of vascular abnormalities marked by CD31 (Figure S13B), suggesting that these macrophages may represent a secondary response to tissue injury and vascular defects in the KO labyrinth (Figure 6E)^61^.

**Figure 6.**
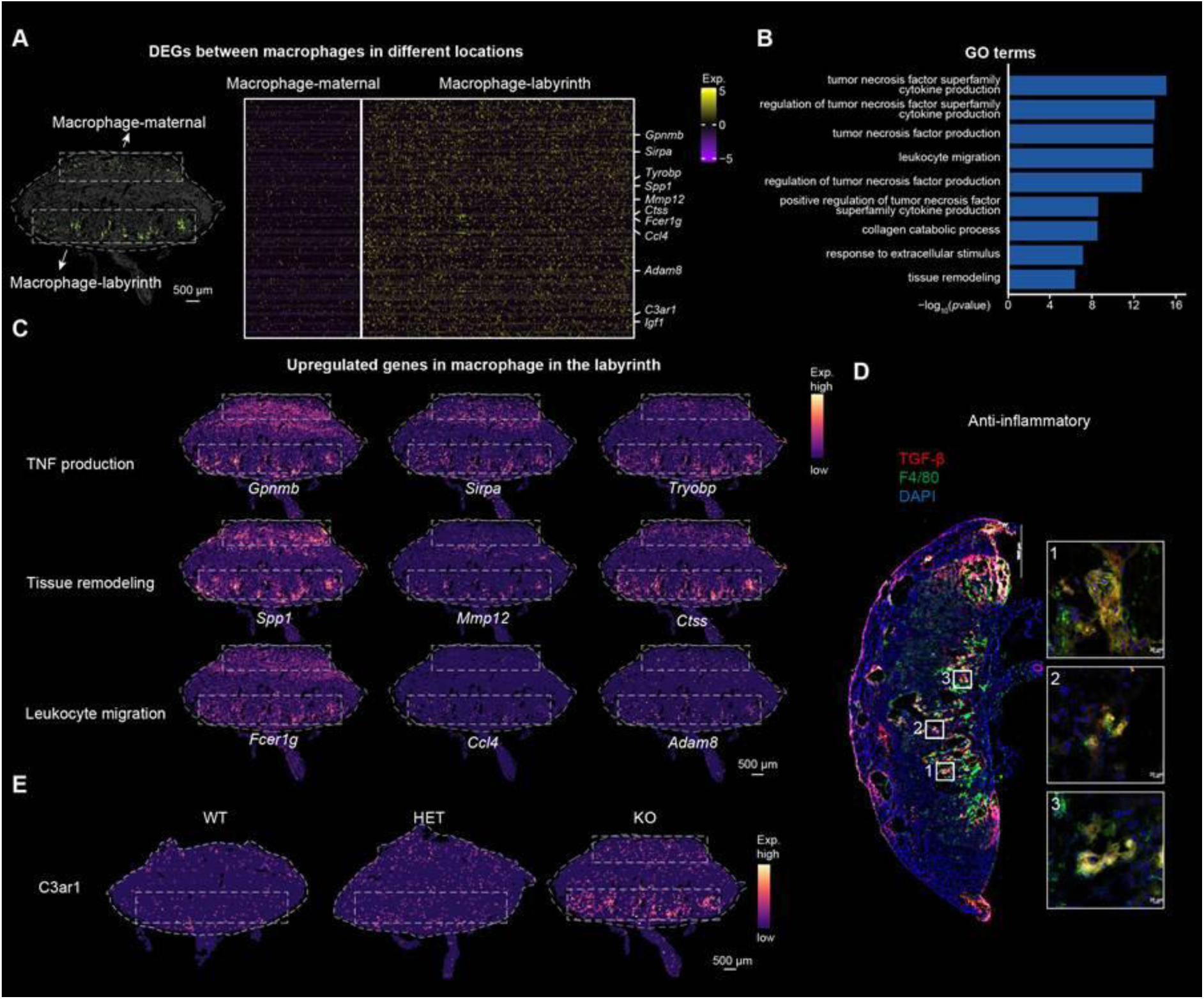
**Secondary macrophage accumulation and immune activation in the labyrinth** (A) Heatmap showing the upregulated genes in macrophages within the labyrinth of KO placenta compared with macrophages within the maternal region. (B) Bar plot showing the enriched GO terms of differentially upregulated genes in macrophages within the labyrinth of KO placentas. (C) Spatial visualization of representative genes in enriched GO terms in the KO placenta section. Scale bars, 500 μm. (D) Co-immunostaining of F4/80 and TGFβ on KO placentas. Scale bars, 1000 μm (overall shape) and 20 μm (high-magnification view). (E) Spatial visualization of *C3ar1* in the WT, HET and KO placenta sections. Scale bars, 500 μm.

## Discussion

The placenta is a complex organ whose proper development is essential for sustaining pregnancy and fetal growth^1,3,62,63,64^. However, how the spatial organization and molecular dynamics of placental cells support this function, and how their disruption contributes to embryonic lethality, remain incompletely understood. Here, we constructed a spatiotemporal transcriptomic atlas (STAMP) of the mouse placenta from E9.5 to E18.5, which delineates cell type composition, spatial organization, and developmental trajectories at single-cell resolution. This atlas revealed a progressive shift of glycogen trophoblast cells (GCs) from the junctional zone (JZ) to the maternal decidua, and uncovered previously unrecognized GC subclusters with distinct molecular profiles and regulatory networks. These findings provide a comprehensive resource for investigating placental ontogeny and pathological perturbations.

Importantly, our study provides direct functional evidence linking placental glycogen metabolism to fetal viability. In a lethal *Ano6*-null model, GCs abnormally persisted at E18.5 and accumulated excessive glycogen granules, while placental and fetal liver tissues exhibited markedly reduced levels of glycogen breakdown intermediates (G1P, G6P, glucose). These results indicate that failure to mobilize placental glycogen deprives the fetus of sufficient metabolic fuel during late gestation, when energy demand peaks^14,16^. Consistently, maternal glucose supplementation restored metabolite levels and significantly improved fetal survival, supporting the notion that GCs serve as an auxiliary energy reservoir to sustain fetal growth under high metabolic demand. This work fills a major knowledge gap by functionally validating the long-suspected role of placental glycogen as a readily mobilizable energy source during late gestation^14,16^.

In addition, our spatial data revealed transcriptional state transitions accompanying GC migration from the JZ to the decidua. GC-1, marked by *Aldh1a3*, arises in the JZ and peaks transiently, while GC-2, marked by *Prl7b1*, emerges after E13.5 in the decidua and expands until E18.5. This spatial shift, validated by RIBO-seq^34^, aligns with classical histological observations of vacuolated, PAS-positive GCs infiltrating the decidua^15,33,65,66^, and highlights the dynamic nature of GC lineage specification. These insights underscore how spatial transcriptomics can capture developmental trajectories and cellular plasticity that are challenging to resolve using dissociative single-cell methods alone.

Beyond the GC-centered metabolic defects, we also observed a secondary phenotype of macrophage accumulation in the labyrinth of *Ano6*-deficient placentas. This feature aligns with previously reported labyrinthine structural defects resulting from impaired trophoblast syncytialization, and likely represents a response to local tissue injury rather than a primary pathogenic mechanism^45^. While not directly implicated in fetal lethality, this finding illustrates the added value of our spatial atlas for uncovering region-specific microenvironmental changes that are invisible to conventional single-cell approaches.

Collectively, our study establishes a spatiotemporal transcriptomic framework for placental development and demonstrates that GC-mediated glycogen metabolism is indispensable for fetal survival. By functionally linking placental glycogen utilization to fetal viability, this work advances our understanding of placental physiology and provides a reference for identifying pathogenic mechanisms underlying pregnancy complications.

## ACKNOWLEDGEMENTS

The authors thank all members of the Liu Lab for their constructive advice and comments on the study. We are grateful to the Imaging Platform at Westlake University for the use of microscopes and to the Histochemical Platform for providing technological support in tissue handling. We would like to thank the Microscopy Core Facility (MCF) of Westlake University for our electron microscopy work and the technical assistance from Yilin Sun and Guicun Fang. We also thank the Laboratory Animal Research Center (LARC) at Westlake University for animal care and operational assistance. Furthermore, we would like to thank CHI BIOTECH CO., LTD. for single nuclei RNA sequencing support. This work was supported by the National Natural Science Foundation of China (32370784, 22DAA01467), the National Key R&D Program of China (2022YFA1105700, 2022YFC3400400) and the Westlake Education Foundation. Additionally, we thank the China National GeneBank for providing sequencing services for this project.

## AUTHOR CONTRIBUTIONS

X.L. conceived the project. X.W., L.L. and X.L. supervised the project. Y.F., S.J., performed all experiments and prepared samples for Stereo-seq and snRNA-seq with help from Y.M., Y.Y., S.W., and X.L. Y.F., X.Z., Y.L. analyzed the data with support from Y.J., T.X., J.S., L.M., X.M., Y.Y., W.P., and L.T. C.Y. and S.S. performed the RIBOmap experiment. Y.F., J.T., M.P.M., S.C., X.W., L.L., and X.L. wrote and revised the manuscript with input from all authors. All authors approved of and contributed to the final version of the manuscript.

## STAR★METHODS

### KEY RESOURCES TABLE

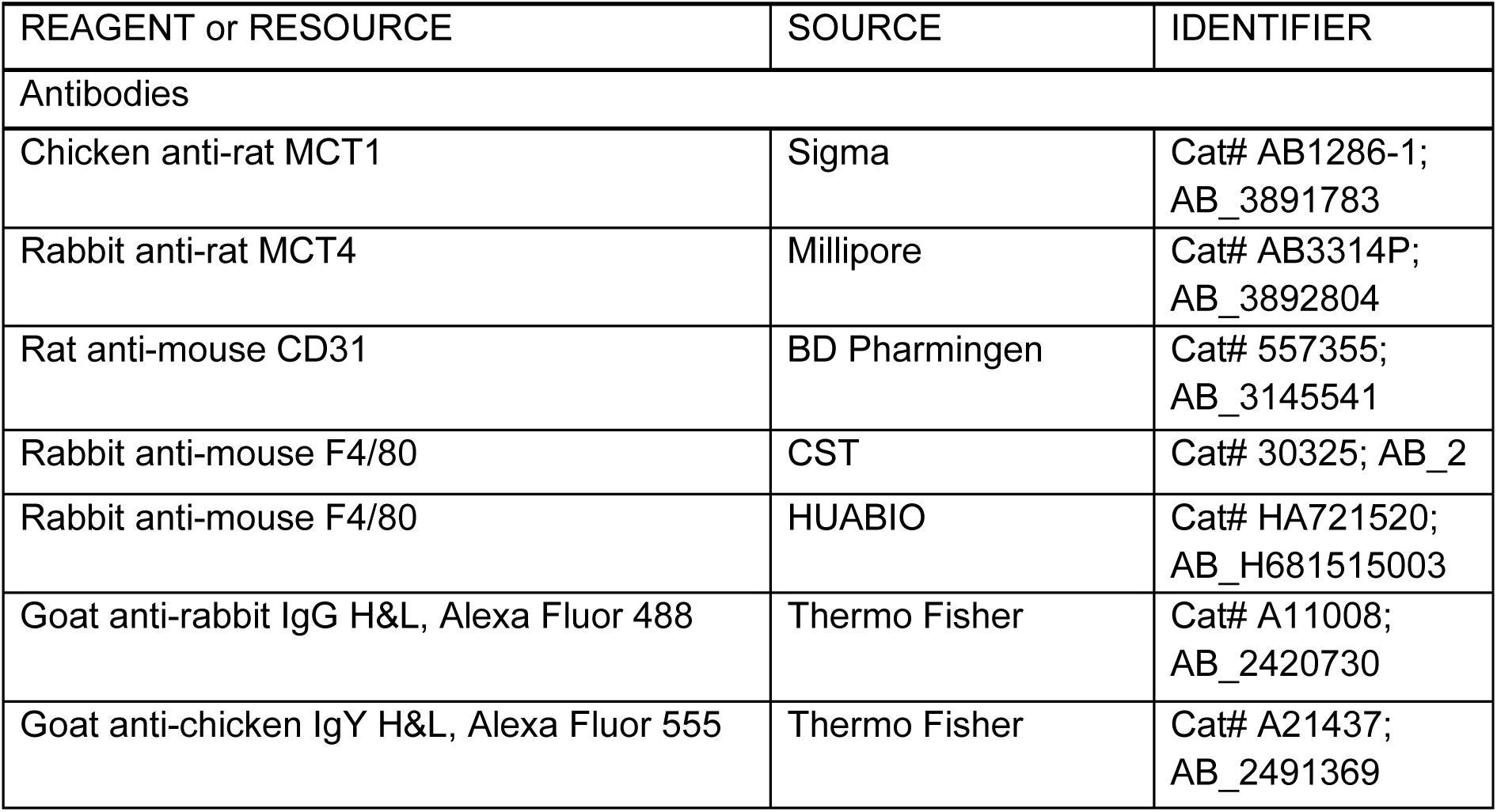
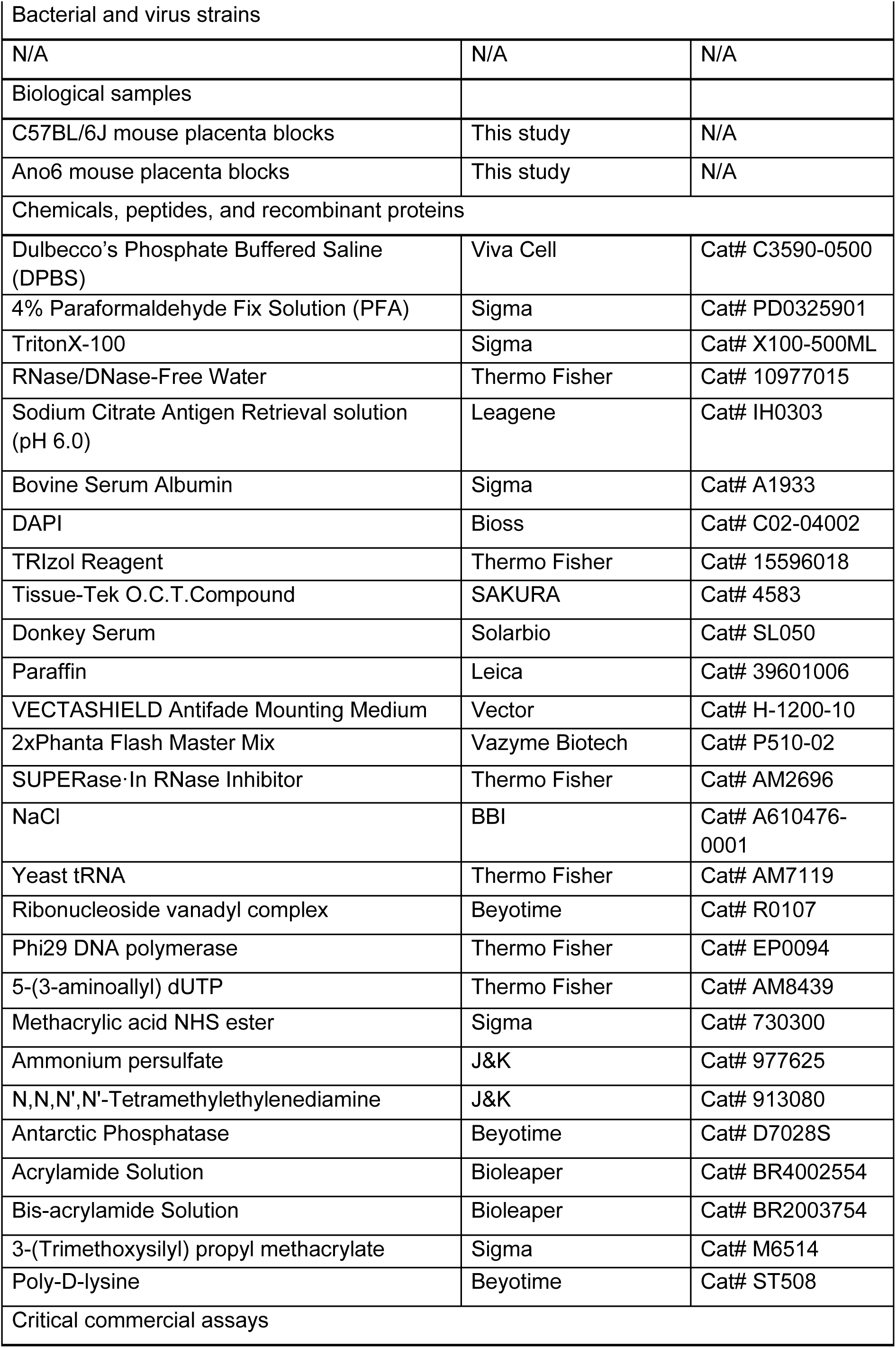
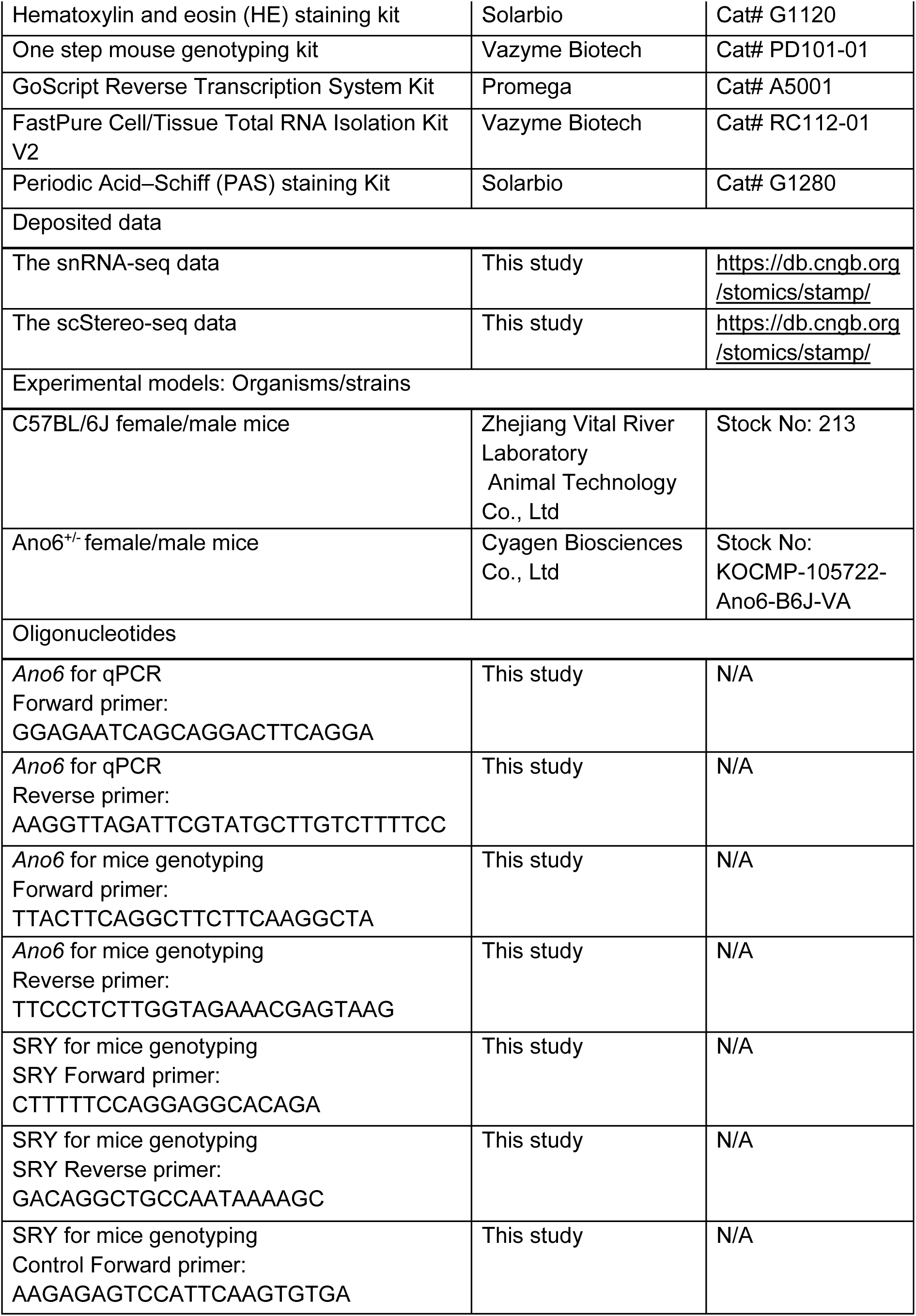
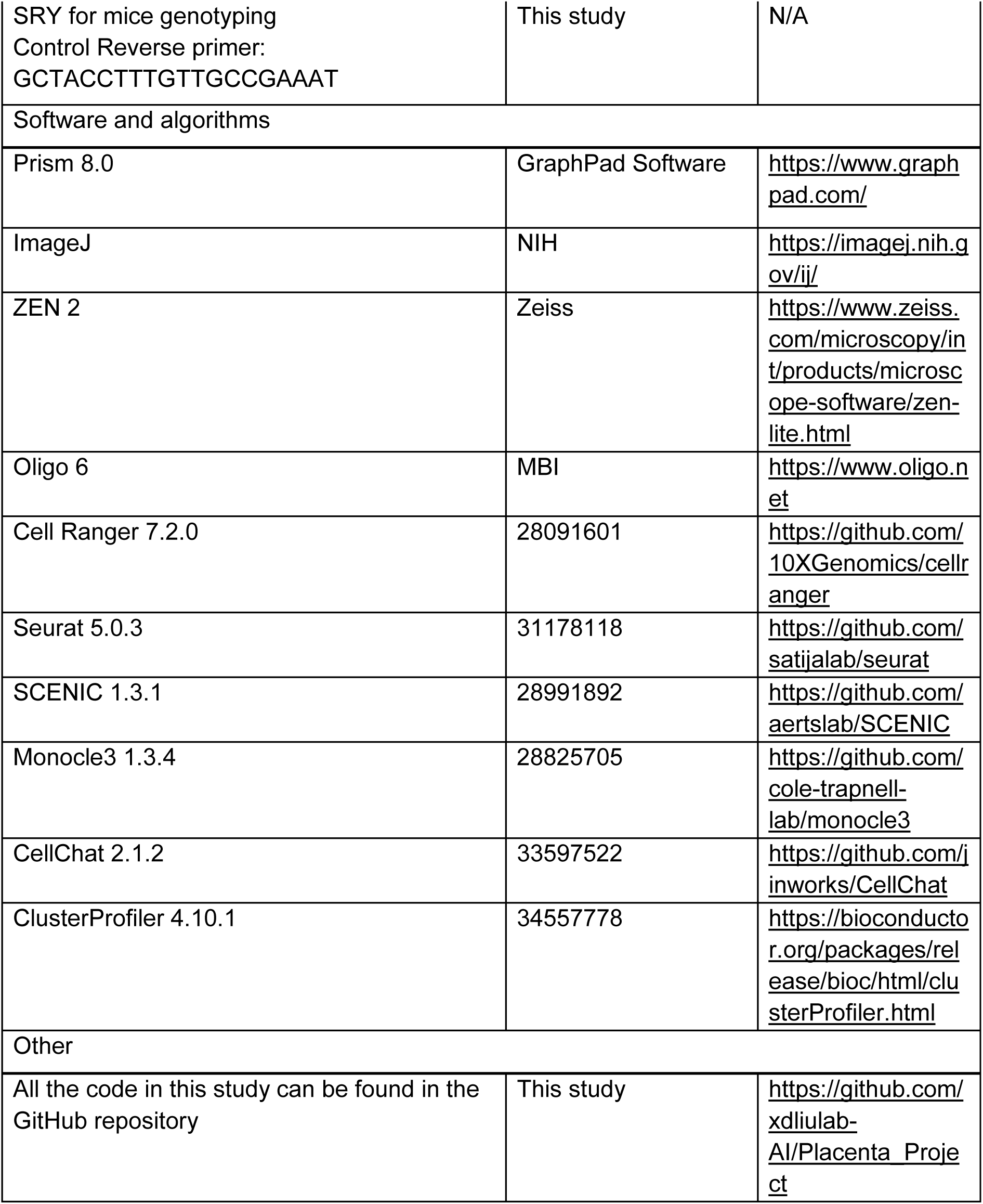

### RESOURCE AVAILABILITY

#### Lead contact

Further information and requests for the resources and reagents may be directed to the corresponding author Xiaodong Liu.

#### Materials availability

This study did not generate new unique reagents. All the reagents in this study were included in the key resources table.

#### Data and code availability

● The data that support the findings of this study have been deposited into CNGB Sequence Archive (CNSA) of China National GeneBank DataBase (CNGBdb) with accession number CNP0004934. The snRNA-seq data and scStereo-seq data are accessible on our interactive data portal at CNSA https://db.cngb.org/stomics/stamp/.
● All the code in this study can be found in the GitHub repository: https://github.com/xdliulab-AI/Placenta_Project.

### EXPERIMENTAL MODEL AND STUDY PARTICIPANT DETAILS

#### Animals

All strains of mice concerned in the study were in the C57BL/6J genetic background. The *Ano6*^+/-^ mouse was generated by Cyagen Biosciences. The knockout (KO) mouse for the *Ano6* gene was generated using a CRISPR/Cas9-mediated strategy. Two guide RNAs (gRNA-B1 and gRNA-B2) were designed to target sequences flanking exon 3 of *Ano6*. Cas9-induced double-strand breaks at these sites resulted in the deletion of the intervening exon, creating a frameshift and functional knockout of the gene. The specific gRNA sequences used were: gRNA-B1: CGGTGACTGCAGTGTTAAGGTTGG; gRNA-B2: CAGAAATCAGCCAAACGAAGCAGG. The *Ano6* mouse mating strategy was crossing heterozygous male mice with heterozygous female mice in order to generate wildtype, heterozygous and homozygous offspring. All animal experiments were carried out with permission of the ethics committee on laboratory animal welfare of Westlake University (AP#22-032-2-LXD-7). All mice were housed in an SPF facility under a 12/12 hours light/dark cycle. Animals were mated at 16:00–17:00, and the vaginal plugs were checked by visual inspection the next morning (8:00–9:00).

### METHOD DETAILS

#### Placenta dissection

Pregnant mice were euthanized, and their placentas were quickly dissected from the uterus. Cold DPBS was used to wash the placentas three times to remove blood. Yolk sac DNA was extracted to confirm genotypes (Vazyme, PD101-01). Surface water was carefully wiped away before the tissue was embedded in OCT (SAKURA, 4583), ensuring that air bubbles were avoided. The frozen blocks were then stored at −80°C. For paraffin embedding, the placentas were fixed overnight in 4% PFA at 4°C, dehydrated through a series of ethanol washes, cleaned three times in xylene, and finally embedded in paraffin (Leica, 39601006) overnight.

#### Stereo-seq experiment

Frozen sections with a thickness of 10 μm were attached to the surface of a 1 cm x 1 cm stereo-seq chip. The tissues were fixed with pre-chilled methanol at −20°C for 30 minutes, then stained with Qubit ssDNA (Invitrogen, Q10212). Tissue permeabilization was performed using STOmics PR Enzyme (1000028500) for 5 minutes. Reverse transcription was conducted at 42°C using the STOmics Gene Expression kit. Bead-purified cDNA was then amplified by PCR, followed by another round of bead purification. For library construction, the cDNA was fragmented and amplified. The products were subsequently filtered twice, first with 0.6X and then with 0.2X beads.

#### Nuclei extraction and snRNA-seq

We prepared 200 mg of frozen tissue and transferred it into a 1.5 ml microcentrifuge tube, taking care not to thaw the tissue before lysis. We added 300 µl of NP40 lysis buffer, minced the tissue using scissors, then added an additional 1 ml of NP40 lysis buffer. We incubated the mixture on ice for 5-8 minutes to allow for lysis. After lysis, the suspension was passed through a 30 µm filter into a 5 ml tube and centrifuged at 4°C at 500 g for 5 minutes. Next, we added 1 ml of PBS containing 1% BSA and 1 U/µl RNase inhibitor, mixed gently by pipetting, and conducted microscopic quality checks. We assessed the integrity of the nuclear membrane and background using Trypan blue under a microscope and applied AOPI stain to evaluate nuclear concentration, viability, and aggregation of nuclei. After confirming satisfactory quality, we proceeded with labeling for downstream analysis as soon as possible.

#### Tissue sectioning

All frozen blocks were equilibrated in the box for at least 30 minutes before sectioning. The frozen embedded tissue was sectioned sagittally until reaching the umbilical cord, with each section being 10 µm thick. Paraffin-embedded sections were cut at a thickness of 5 µm. For the histological examination of the placenta, at least three sections, positioned 70 µm apart, were analyzed per placenta.

#### Hematoxylin-Eosin (H&E) staining

Frozen slices were allowed to recover for 20 minutes at room temperature before being fixed in 4% PFA for 3 minutes. Tissue sections were stained with hematoxylin for 6 minutes and with eosin for 2 minutes. Dehydration was carried out through an alcohol gradient of 75%, 85%, 95%, and 100%, followed by two xylene washes, each lasting 1 minute. Paraffin slices were incubated in hematoxylin for 6 minutes, rinsed in water, and then stained in eosin for 30 seconds. Spatial transcriptomic sections and all validation analyses (H&E, IF, and RIBOseq) were prepared from adjacent or same-block placental regions to ensure anatomical comparability across datasets.

#### Immunofluorescence staining

Placentas were dissected and post-fixed in 4% PFA/PBS for 30 minutes, then permeabilized in 0.1% Triton X-100/PBS for 30 minutes. Blocking was performed with 5% donkey serum and 1% BSA for 30 minutes at room temperature. Sections were incubated with diluted primary antibodies overnight at 4°C, washed three times with PBS, and then incubated with secondary antibodies for 1 hour at room temperature. Primary and secondary antibodies were used at the following concentrations: chicken anti-rat MCT1 (sigma, AB 1286-1, 1:200), rabbit anti-rat MCT4 (Millipore, AB3314P, 1:100), rat anti-mouse CD31 (BD Pharmingen, 557355, 1:200), rabbit anti-mouse F4/80 (CST, 30325, 1:200), rabbit anti-mouse F4/80 (HUABIO, HA721520, 1:100), goat anti-rabbit IgG H&L (Abcam, Alexa Fluor 488, 1:500), goat anti-chicken IgY H&L (Abcam, Alexa Fluor 555, 1:500).

#### Periodic Acid–Schiff (PAS) staining

PAS staining was performed using a PAS staining kit (Cat #G1280, Solarbio) following the manufacturer’s instructions. Paraffin sections were dewaxed to distilled water. The sections were treated with an oxidant for 10 minutes and subsequently rinsed in running tap water for 5 minutes. Samples were then stained with Schiff reagent for 15 minutes and rinsed again for 5 minutes. The sections were counterstained with Mayer’s hematoxylin solution for 3 minutes. Following conventional dehydration and clearing in xylene, the sections were mounted with neutral gum. Quantification of the PAS-positive area relative to the total placental area for each section was performed using ImageJ software.

#### Quantitative PCR (qPCR)

Total RNA was extracted using the FastPure Cell/Tissue Total RNA Isolation Kit V2 (Vazyme RC112). Each RNA sample was reverse-transcribed with the GoScript Reverse Transcription system (A5001, Promega) using Oligo(dT) and random primers. Relative quantitation was determined using the BioRad CFX Touch and calculated by the comparative Ct method (2−ΔΔCt), with the expression of β-Actin serving as the control.

#### Ribosome-bound mRNA mapping (RIBOmap)

10 μm thick placenta slices were mounted on confocal dishes. Tissues were fixed in 4% PFA for 15 minutes and permeabilized with cold methanol at −20°C for 1 hour. Yeast tRNAs were added to quench the tissues. Splint primers, padlock, and primer probes were added to the tissue and hybridized for 12 hours at 40°C. After washing twice in PBSTR and once with a high-salt washing buffer for 20 minutes each at 37°C, T4 ligase was added to the solution and incubated for 2 hours at room temperature. Rolling circle amplification of the DNA circle by phi29 polymerase was performed at 34°C for 30 minutes and at 30°C for 2 hours. Tissues were washed twice with 0.1% PBST. After incubation in modification solution for one hour at room temperature, tissues were monomerized with 4% acrylamide-0.2% bis-acrylamide for 15 minutes. Coverslips were then placed directly onto the polymerization solution and incubated under nitrogen flow for 1 hour, after which the coverslip was discarded. Tissues were digested in PK buffer mix (0.2 mg/ml PK final) for 1 hour at 37°C. Placentas were stripped for 10 minutes in 60% formamide twice. After 3 hours of incubation at room temperature, the fluorescent oligos were visualized by confocal fluorescence microscopy using LAS X software.

#### Transmission electron microscopy (TEM)

Placental tissues were rapidly collected and immersed in a solution of 2.5% glutaraldehyde and 2% paraformaldehyde in 0.1 M phosphate buffer (PB) at 4°C overnight for fixation. Post-fixation, the samples were washed three times with PB buffer and subsequently fixed with 2% osmium tetroxide for 60 minutes on ice. This was followed by a further fixation in 2% osmium tetroxide containing 2.5% potassium ferrocyanide for an additional 60 minutes on ice. The samples were dehydrated through a graded ethanol series (30%, 50%, 70%, 85%, 95%, and twice in 100%) for 10 minutes each step. They were then infiltrated and embedded in EPON12 resin. After sectioning into ultra thin slices (70 nm) and staining, the sections were imaged using a Thermo Fisher Talos 120 electron microscope operated at 80 kV. Quantification of glycogen granules per μm² in glycogen cells was performed using ImageJ software.

#### Placental glycogen content quantification

Whole placentas were immediately weighed on a scale. Each sample was treated with 0.5 mL of 30% KOH saturated with Na_2_SO_4_. The tubes were then placed in a boiling water bath for 30 minutes until a homogeneous solution formed. After removal from the bath, the tubes were cooled on ice. To precipitate the glycogen, 95% ethanol was added. The samples were left to stand on ice for 30 minutes, followed by centrifugation at 840 g for 30 minutes. The glycogen precipitates were dissolved in 3 mL of distilled water. To each mL of sample, 1 mL of 5% phenol solution was added, followed by the rapid addition of 5 mL of 98% H_2_SO_4_. The tubes were allowed to stand for 10 minutes, shaken, and then incubated at 30°C for 20 minutes. Blanks were prepared using 1 mL of distilled water. Absorbance was measured at 490 nm. Quantification of glycogen content and the percentage of glycogen weight relative to total placental weight were performed using GraphPad Prism software.

### QUANTIFICATION AND STATISTICAL ANALYSIS

#### Data Processing of snRNA-seq Data

Single-nucleus RNA sequencing data processing utilized Cell Ranger (version 7.2.0, 10x Genomics) for quality control, alignment to the Mus musculus genome (refdata-gex-mm10-2020-A), and transcript quantification. The integration step employed the Cell Ranger aggr pipeline for sequencing depth normalization. Post-integration, cells with fewer than 500 genes detected or over 10% mitochondrial expression were excluded. The remaining data underwent normalization with Seurat’s NormalizeData function (version 5.0.3).

#### Batch Correction and UMAP Nonlinear Dimensionality Reduction

The dataset underwent batch correction and dimensional reduction using IntegrateLayers with FastMNNIntegration. Neighborhood identification uses FindNeighbors (reduction = ’integrated.mnn’, dims = 1:30), followed by cluster identification with FindClusters (resolution = seq(0.1, 0.2, 0.1)). Dimensional reduction concluded with RunUMAP (dims = 1:30, reduction = ’integrated.mnn’), visualizing cell clusters in two dimensions.

#### Identification of Differentially Expressed Genes

Differential gene expression was determined using FindMarkers, focusing on genes with only positive expression in specific clusters, detected in at least 25% of cells (min.pct = 0.25) and showing significant expression differences (logfc.threshold = 0.25).

#### SCENIC Analysis

Seurat objects converted to loom format enabled SCENIC analysis. GRNBoost2 from Arboreto inferred gene regulatory networks. The workflow involved runSCENIC_1_coexNetwork2modules, runSCENIC_2_createRegulons, and runSCENIC_3_scoreCells, assessing regulon activity across cell types with cisTarget databases set to mm10, and visualized through heatmaps.

#### Pseudotime Analyses and Characterization of GC Subtypes

Monocle 3 was used to transform Seurat objects into cell datasets and cluster cells (resolution = 1e-3). Trajectory analysis with learn_graph (use_partition = TRUE) identified developmental pathways, with order_cells setting the JZP cluster as the starting point. Visualization focused on pseudotime and cluster trajectories.

#### Cell-Cell Communication Analyses

Using CellChat, cell-cell communication networks were analyzed, utilizing the CellChatDB mouse database. IdentifyOverExpressedGenes and identifyOverExpressedInteractions pinpointed crucial genes and interactions. computeCommunProb (type = “triMean”) and filterCommunication (min.cells = 10) refined the communication data.

#### Cell Type Integration

Data integration across sources standardized Seurat objects to the “RNA” assay using IntegrateLayers and “FastMNNIntegration”. Neighborhoods were defined with FindNeighbors (dims = 1:30), and clusters identified with FindClusters. Visualization was achieved with RunUMAP, harmonizing and analyzing integrated datasets.

#### Ontology Annotation

GO enrichment analysis with clusterProfiler and org.Mm.eg.db mapped mouse genes to GO terms. Gene symbols converted to Entrez IDs with bitr preceded enrichGO analysis for clusters (snRNA-seq: pvalueCutoff = 0.01, pAdjustMethod = “fdr’’, minGSSize = 10, maxGSSize = 500, qvalueCutoff = 0.01; Stereo-seq: pvalueCutoff = 0.05, pAdjustMethod = “BH’’, minGSSize = 10, maxGSSize = 500, qvalueCutoff = 0.2).

#### Stereo-seq raw data processing

Single-end Stereo-seq fastq files were generated using a MGI DNBSEQ-Tx sequencer, which contained CID (coordinate identity), MID (molecular identity, UMI) and cDNA sequences. The retained reads were aligned to the mouse reference genome (mm10) (https://ftp.ensembl.org/pub/release-93/fasta/mus_musculus/dna/Mus_musculus.GRCm38.dna.primary_assembly.fa.gz) by STAR. Only mapped reads with a mapping quality score (MAPQ) > 10 were then annotated and calculated using handleBam (available at https://github.com/BGIResearch/handleBam). UMIs with identical CID and gene locus were collapsed, allowing 1 base pair of mismatch to correct for sequencing and PCR errors. Exonic reads were then used to construct an expression profile matrix comprising CIDs.

#### Image-based Cell Segmentation of Stereo-seq data

We utilized ssDNA staining image from the same section to segment cells by projecting the ssDNA staining image onto the Stereo-seq chip image. Firstly, a grayscale map was generated from the Stereo-seq data, with each pixel representing one DNB. Manual registration was conducted on both the grayscale map and the ssDNA staining image to align the pixels of the staining image with the DNB coordinates. Cell segmentation was then carried out using the scikit-image python package (v0.19.3)^67,68^. Global threshold was used to filter the background noise of the registered image, and a mask was generated for segmentation. Gaussian-weighted local threshold was calculated to represent the position of the cell with block size of 45 and offset of 0.03. Cells with overlapped regions were segmented using the exact Euclidean distance transformation (with a minimal distance of 15). We then expanded the labels representing different cells in the label image by 5 pixels without causing overlaps, via expand_labels function. For each segmented cell, UMIs from all DNBs within the corresponding segmentation were aggregated per gene and subsequently summed to generate a cell-by-gene matrix for downstream analysis.

#### Spatially-constrained clustering of Stereo-seq data and region annotation

Cells with UMI counts below 200 (100 for E9.5, E15.5) were excluded from further analysis. The remaining raw count matrices of the placenta samples were normalized by SCTransform function in Seurat (v4.3.0)^69^. Spatial information was taken into account for unsupervised clustering. The centroid of each cell was computed based on its spatial coordinates. The spatial k-nearest neighbor graph 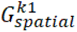 was constructed using Squidpy (v1.2.3)^70^. This graph was then combined with the k-nearest neighbor graph 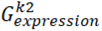 based on transcriptomic data (*k*_2_ is by default set to be 30). The combined graph 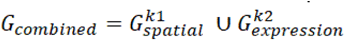 was used as input for Leiden clustering. We annotated the placental region according to the anatomical structure, combined with spatially-constrained clustering and HE section partitioning.

#### Single cell level annotation of Stereo-seq data

Cells with UMI counts below 200 (100 for E9.5, E15.5) were excluded from further analysis. The remaining raw count data were normalized by the SCTransform function in Seurat (v4.3.0) to mitigate the effects of sequencing depth. Subsequently, high-quality single-nucleus RNA sequencing (snRNA-seq) data from the corresponding developmental stage were employed as a reference to annotate the Stereo-seq data using the TACCO (v0.3.0) framework^26^. Specifically, we mapped the Stereo-seq data using snRNA-seq data as a reference and the adjusted snRNA-seq data cell type proportions as the annotation prior distribution via the tacco.tools.annotate function. The annotation process assigned each spatial single cell the cell type with the highest score, indicating the most probable identity.

## Additional Information

### Supplementary figure

**Figure S1.**
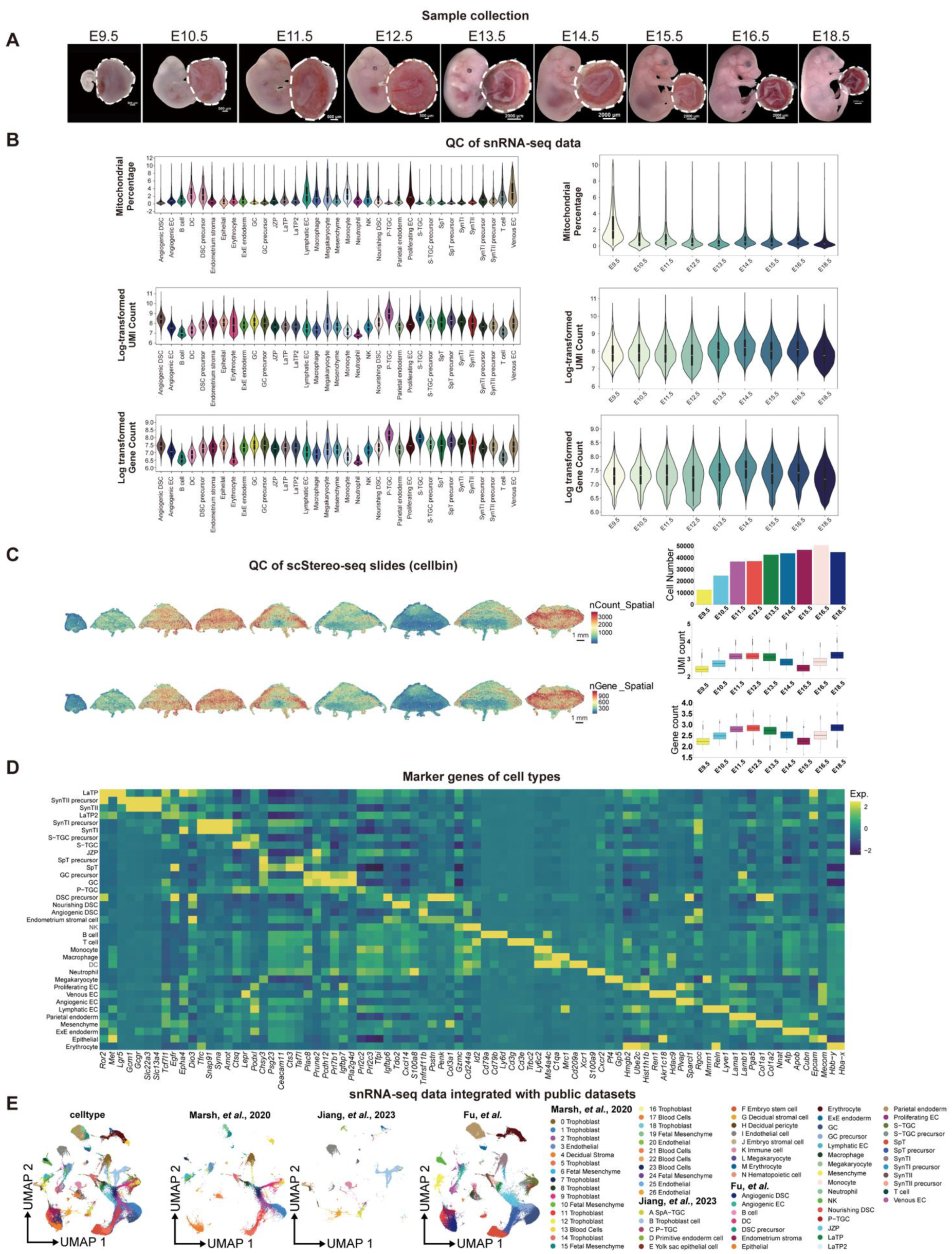
Experimental design and quality control analysis of snRNA-seq and Stereo-seq dataset, related to. **Figure 1** (A) Bright field images of samples from different developmental stages collected for sequencing experiments. White dashed lines indicate the collected placenta samples. (B) Violin plots showing the number of reads and genes, and percentage of mitochondrial genes in the snRNA-seq data. (C) Quality control of Stereo-seq data on cell number, UMI count and gene count. Stereo-seq spot overlay (cellbin) showing number of reads and genes. Scale bars, 500 μm. (D) Heatmap showing top gene expression of all identified cell types. (E) Integration of snRNA-seq data in this study with published datasets.

**Figure S2.**
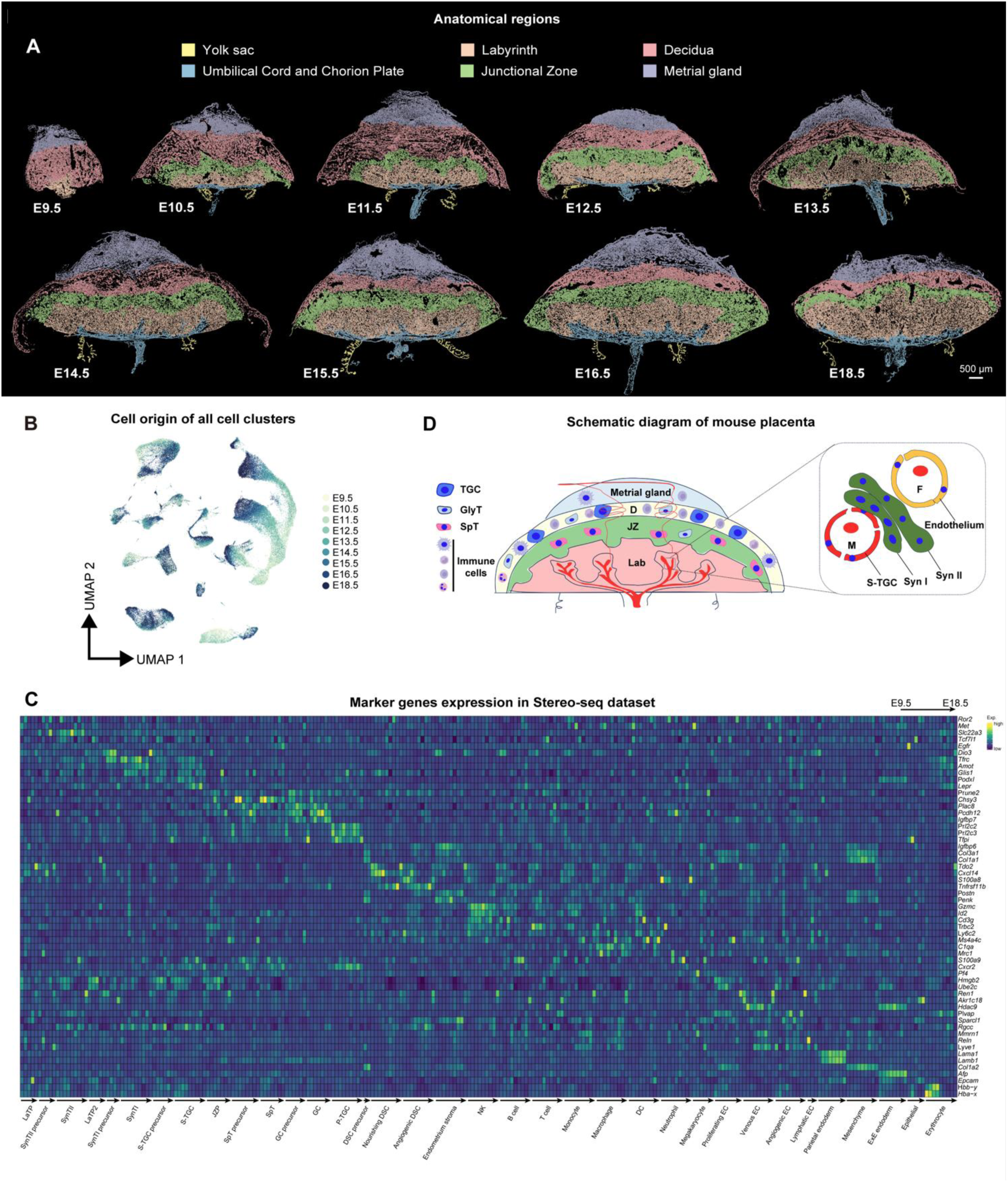
Anatomical regions and marker gene expression in Stereo-seq, related to. **Figure 1** (A) Anatomical regions were identified. Regions were colored based on anatomical region annotation. Scale bars, 500 μm. (B) UMAP displaying snRNA-seq data from different time points. (C) Heatmap showing marker gene expression of all identified cell types using Stereo-seq dataset. (D) Schematic diagram of annotated placental regions based on our scStereto-seq data analysis.

**Figure S3.**
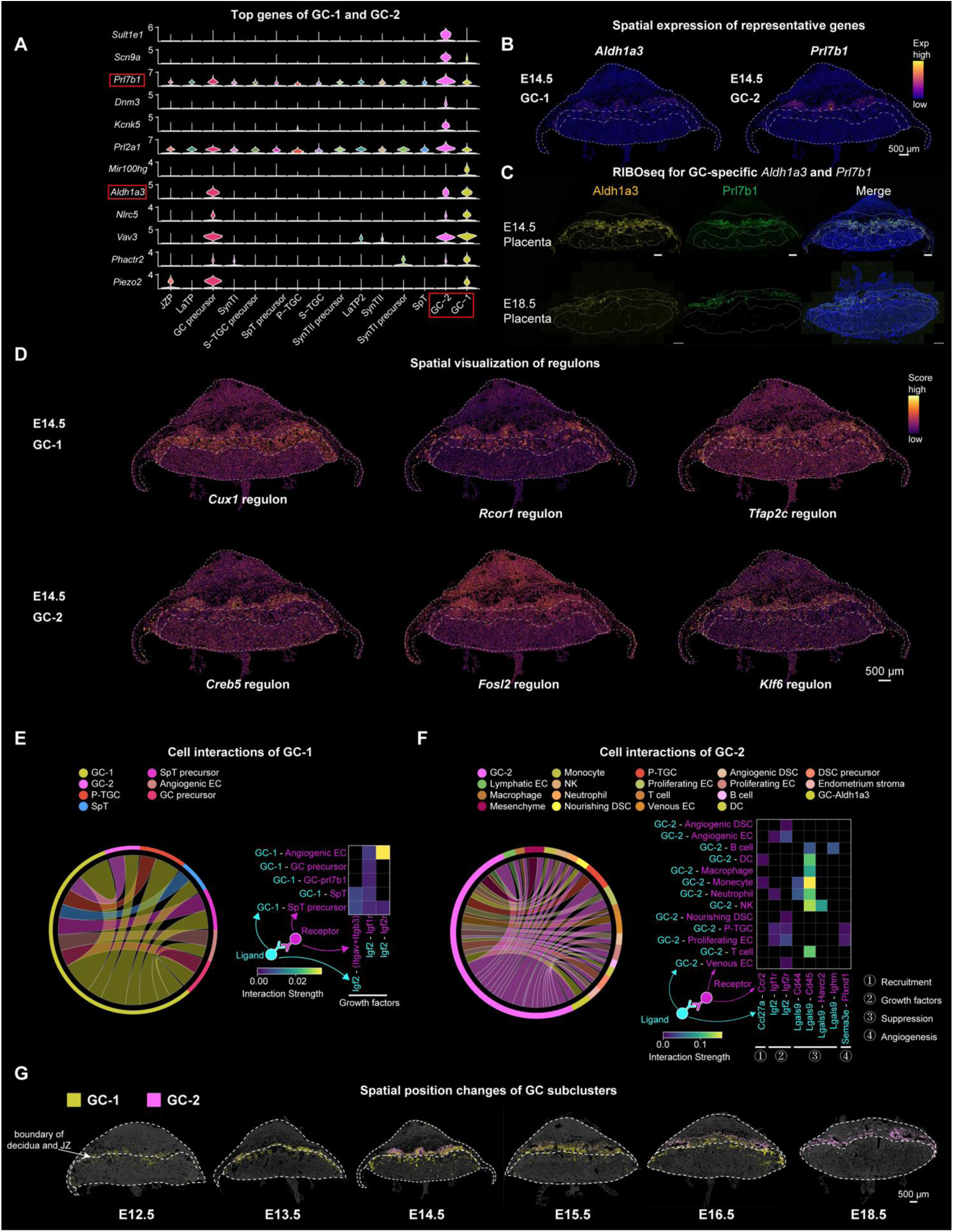
Molecular differences between two GC subclusters, related to. **Figure 2** (A) Violin plot showing top genes of GC-1 and GC-2 in all trophoblast cells. (B) Spatial expression of top genes, *Aldh1a3* and *Prl7b1*, in GC-1 and GC-2 on an E14.5 placental section, respectively. Scale bars, 500 μm. Dotted lines encircle the regional boundaries. Scale bars, 500 μm. (C) Detection of *Aldh1a3* and *Prl7b1* protein synthesis by RIBOseq in E14.5 and E18.5 placental sections. Dotted lines encircle the regional boundaries. Scale bars, 500 μm. (D) Spatial visualization of selected regulons in GC-1 and GC-2 on E14.5 placental sections, respectively. (E) Chord (left) diagram showing the interactions between GC-1 (sender) and other cell types (receiver). Heatmap (right) showing the selected ligand-receptor interactions. (F) Chord (left) diagram showing the interactions between GC-2 (sender) and other cell types (receiver). Heatmap (right) showing the selected ligand-receptor interactions. (G) The spatial visualization of the two GC subclusters from E12.5 to E18.5. Cells are colored by their annotation. Scale bars, 500 μm. Inner dotted lines represent the boundary of decidua and JZ. Scale bars, 500 μm.

**Figure S4.**
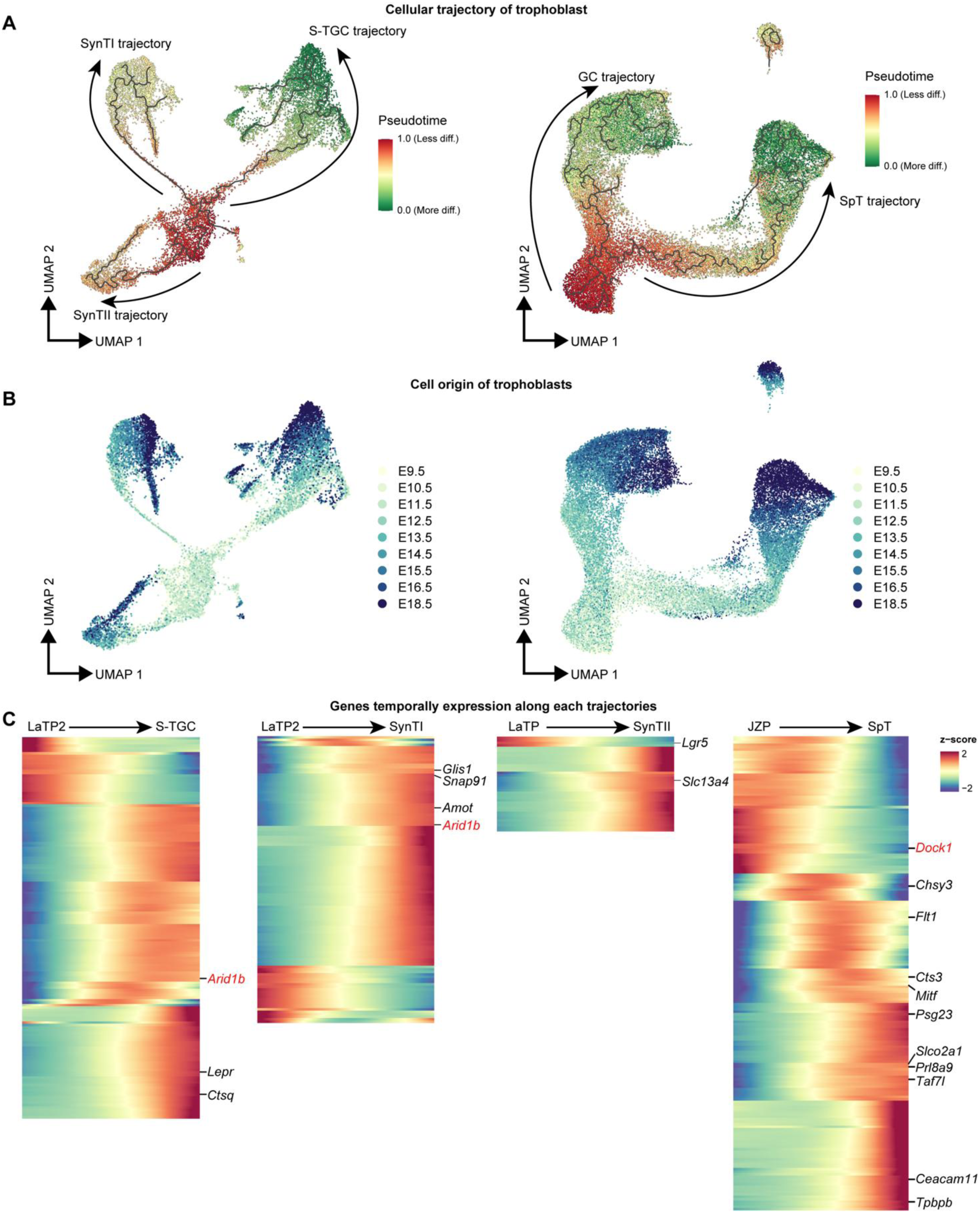
Cellular trajectory analysis identifies genes with temporally regulated expression patterns along pseudotime, related to. **Figure 3** (A) Cellular trajectory reconstruction of trophoblast cells using the Monocle3 and CytoTRACE. (B) UMAP visualization showing the origin of trophoblasts based on the time point. (C) Z-score heatmap of gene expression in different branches, where rows are genes and columns are cells ranked by pseudotime value. Genes were first fit with Moran’s I test with ranked pseudotime as independent variable. Genes with the most significant time dependent model (*q*_value=0 and morans_I>0.25) were extracted and clustered hierarchical clustering. Cells were ordered according to scaled pseudotime value from 0 to 1. Genes previously identified to be associated with lineage development are labeled in black, while potential regulators whose loss leads to embryonic lethality are highlighted in red.

**Figure S5.**
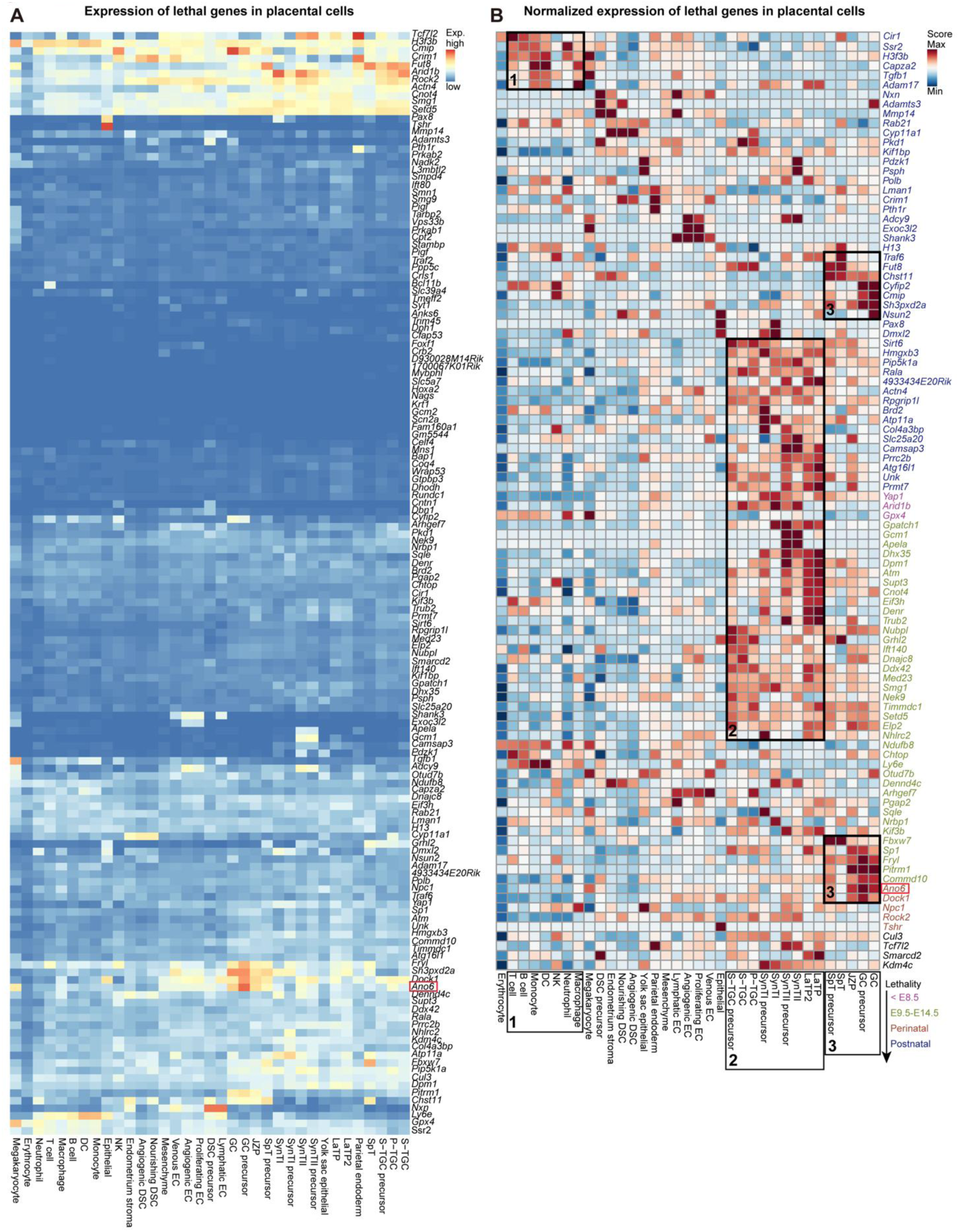
Expression pattern of embryonic lethal genes in placental cells, related to. **Figure 3** (A) Heatmap showing the expression patterns of 151 genes listed in Table 1 in annotated cell types across E9.5-18.5 placentas (snRNA-seq data). The *Ano6* gene is highlighted with a red box. (B) Heatmap showing the normalized expression patterns of top 98 genes selected from Table 1 (based on the expression threshold) in annotated cell types across E9.5-18.5 placentas (snRNA-seq data). The mutant genes are colored with categories by the timing of lethality. The mutants represented in blue are postnatal lethal; the mutants represented in pink are lethal before E8.5; the mutants represented in green are lethal between E9.5-E14.5; the mutants represented in brown are perinatal lethal; and the mutants represented in black are lethal with uncertain embryonic stage. The boxes indicated selected enriched patterns, and the corresponding specific cell types of placentas are also encircled below. The *Ano6* gene is highlighted with a red box.

**Figure S6.**
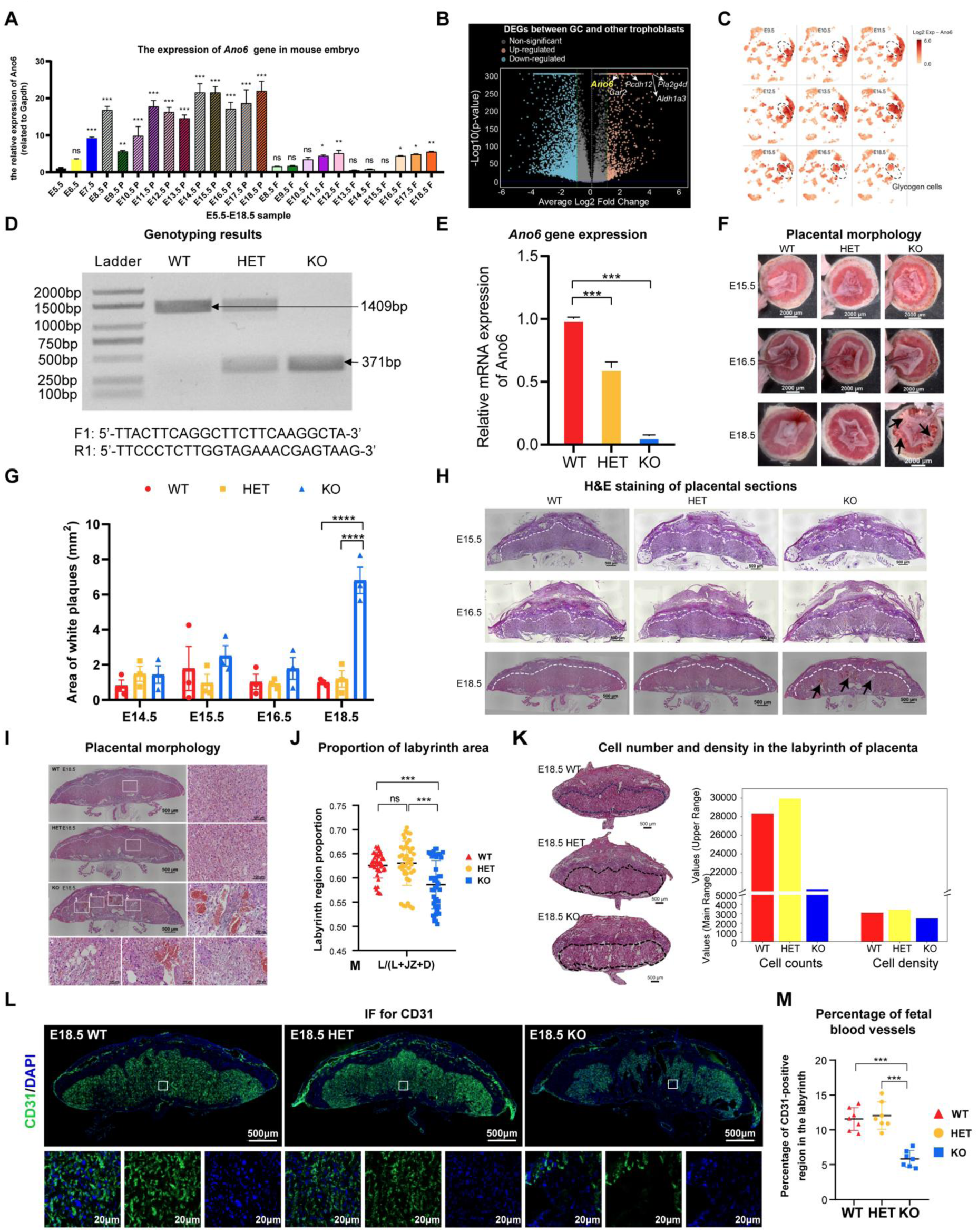
Characterization of *Ano6* WT, HET and KO placentas, related to. **Figure 3** (A) The expression changes of *Ano6* gene in mouse embryos from E5.5 to E18.5 examined using quantitative PCR (qPCR), separately analyzing the expression in the fetal part and the placental part after E8.5. One-way analysis of variance (ANOVA). \**p* < 0.05, \*\**p* < 0.01, \*\*\**p* < 0.001. All data represent means ± SEM. (B) Volcano plot of differentially expressed genes (DEGs) identified between GC and other trophoblast cells. The orange dots denote up-regulated gene expression, the blue dots denote down-regulated gene expression, and the gray dots denote gene expression without marked differences. Several up-regulated genes in GC related to embryonic lethality are highlighted. (C) Ano6 expression in placental cells across different developmental stages, as revealed by snRNA data. The dashed circle highlights the GC lineage. (D) Genotyping results of WT, HET and KO *Ano6* placentas. (E) qPCR analysis of *Ano6* gene in WT/HET/KO placentas at E18.5. One-way analysis of variance (ANOVA). \*\*\**p* < 0.001. All data represent means ± SEM. (F) Representative images of placentas from WT, HET, and KO mice at E15.5, E16.5, and E18.5. Black arrows point to the phenotypic defects in the KO placenta. Scale bars, 2000 μm. (G) Quantification of white plaque area (mm²) on the fetal surface of the placenta across E14.5 to E18.5 in WT, HET, and KO mice. One-way analysis of variance (ANOVA). \*\*\**p* < 0.001. Data are presented as mean ± SEM. (H) H&E staining of WT, HET, and KO placental sections at E15.5, E16.5, and E18.5. The black arrow indicates the location where phenotypic abnormalities appear. Scale bars, 500 μm. The dashed lines indicate the boundary of the labyrinth region. (I) H&E staining of WT/HET/KO placentas at E18.5. Scale bars, 500 μm (overall shape) and 100 μm (high-magnification view). (J) Statistics on the proportion of the labyrinth region in the total area of the placenta. One-way analysis of variance (ANOVA). \*\*\**p* < 0.001. All data represent means ± SEM. (K) H&E staining of WT/HET/KO placentas (left, the labyrinth region was encircled), and statistical results of cell quantity and cell density in the labyrinth region of WT, HET, and KO placentas (right). Cell density was calculated as the number of nuclei divided by the labyrinth area. Scale bars, 500 μm. (L) CD31 immunofluorescence staining of the WT/HET/KO placentas at E18.5. CD31 labels fetal blood vessels. Scale bars, 500 μm (overall shape) and 20 μm (high-magnification view). (M) Percentage of fetal blood vessels in the labyrinth region. WT: *n*=7, HET: *n*=7, KO: *n*=7. One-way analysis of variance (ANOVA). \*\*\**p* < 0.001. All data represent means ± SEM.

**Figure S7.**
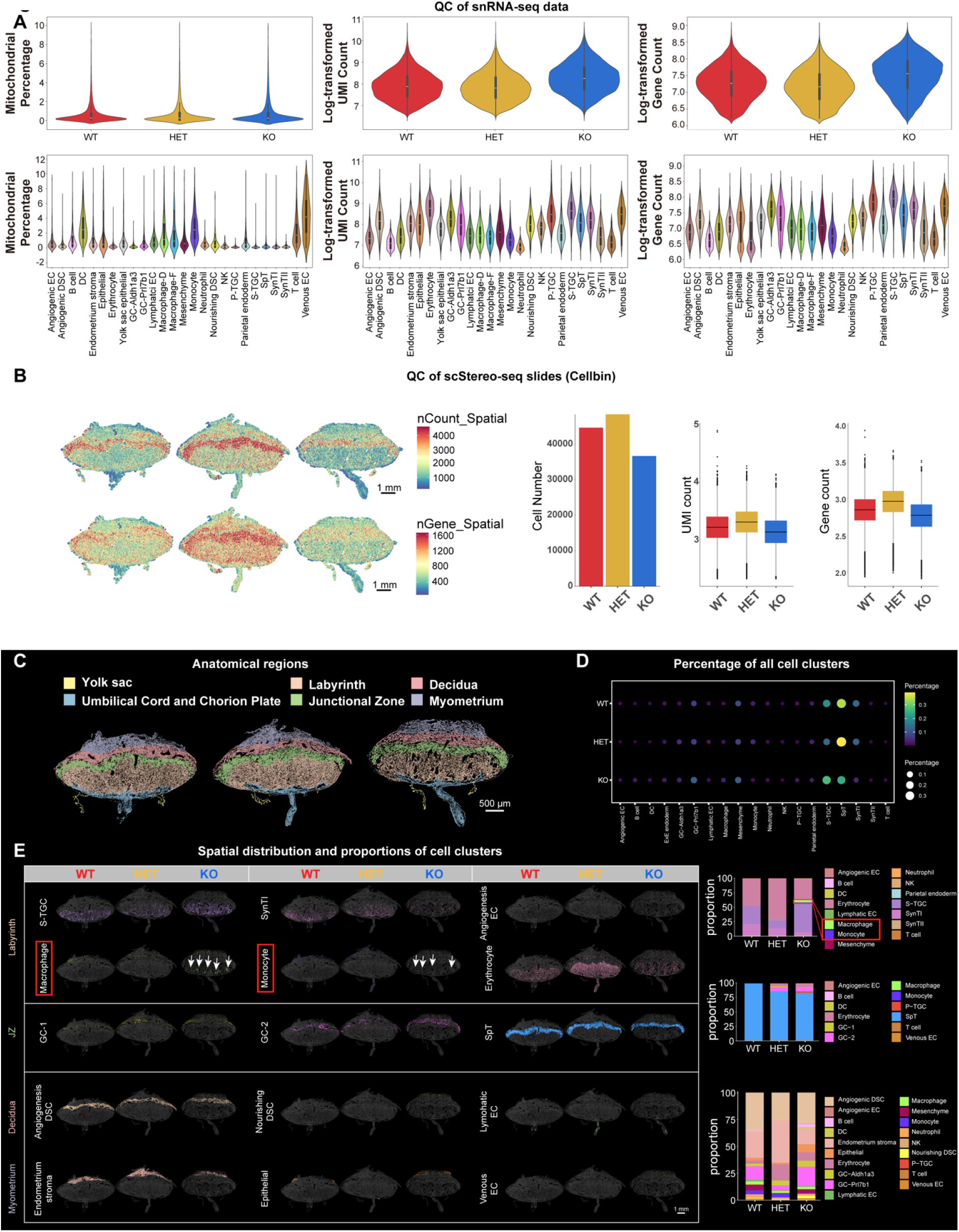
Quality control analysis and characterization of snRNA-seq and Stereo-seq data for E18.5 WT, HET and KO placentas, related to. **Figure 3** (A) Violin plots showing the mitochondrial percentage, UMIs and genes in WT/HET/KO placentas (top) and in each cell type (bottom). (B) Quality check of Stereo-seq data. Stereo-seq spot overlay (cellbin) showing number of genes and reads. Scale bars, 500 μm. (C) Identification of anatomical regions on WT/HET/KO placentas. Regions are colored based on anatomical region annotation. Scale bars, 500 μm. (D) Bubble plot showing the percentage of all cell types in the E18.5 WT, HET, and KO placentas using Stereo-seq data. (E) Cell type distribution in E18.5 WT, HET, and KO placenta sections. Differences in *Ano6* KO placenta compared with the WT/HET placentas are marked red. Cells were colored based on cell type annotation. Bar plots showing cell subcluster proportions in the corresponding regions. Scale bars, 500 μm.

**Figure S8.**
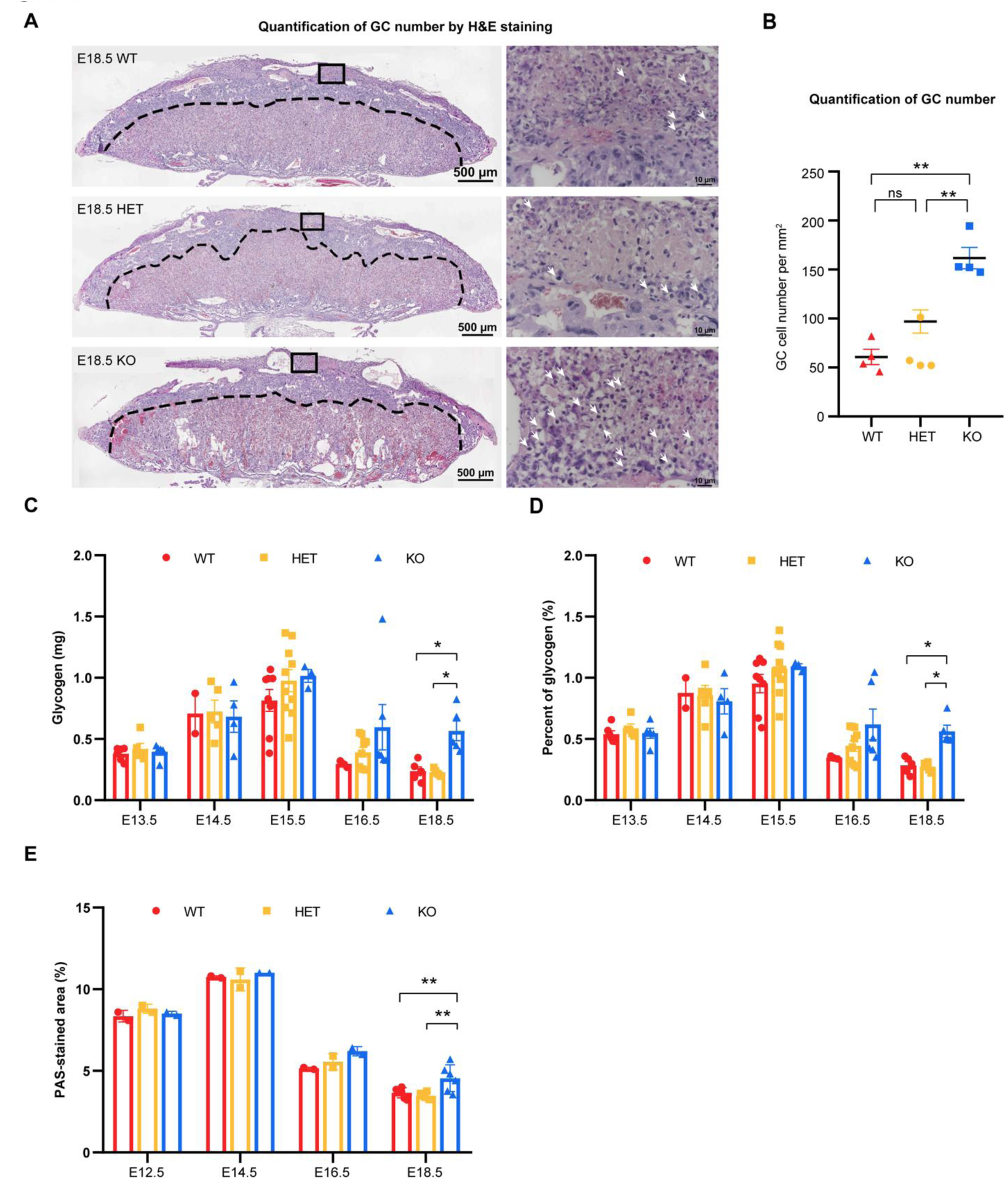
Time-course analysis of placental glycogen content in the defective placenta, related to. **Figure 4** (A) H&E staining of WT, HET and KO placentas. The black dashed line indicates the boundary between the JZ and decidua. The glycogen cells in JZ and decidua are characterized by vacuolated, glycogen-rich cytoplasm and appear as compact cell islets. Scale bars, 500 μm (overall shape) and 10 μm (high-magnification view). Quantification was performed on GCs located in both the junctional zone (JZ) and decidua. (B) The glycogen cell number in JZ and decidua per mm^2^. WT (n=4), HET (n=4), KO (n=4). One-way analysis of variance (ANOVA). \*\**p* < 0.01. All data represent means ± SEM. (C) Total placental glycogen content (mg) in WT, HET, and KO placentas across E13.5 to E18.5. One-way analysis of variance (ANOVA). \**p* < 0.05. Data are presented as mean ± SEM (n = 3-5 per group). (D) Glycogen content expressed as a percentage of total tissue weight (%) in placental tissues across the same developmental stages. \**p* < 0.05. (E) Percentage of PAS-stained positive area relative to the total tissue area analyzed using ImageJ. \*\**p* < 0.01.

**Figure S9.**
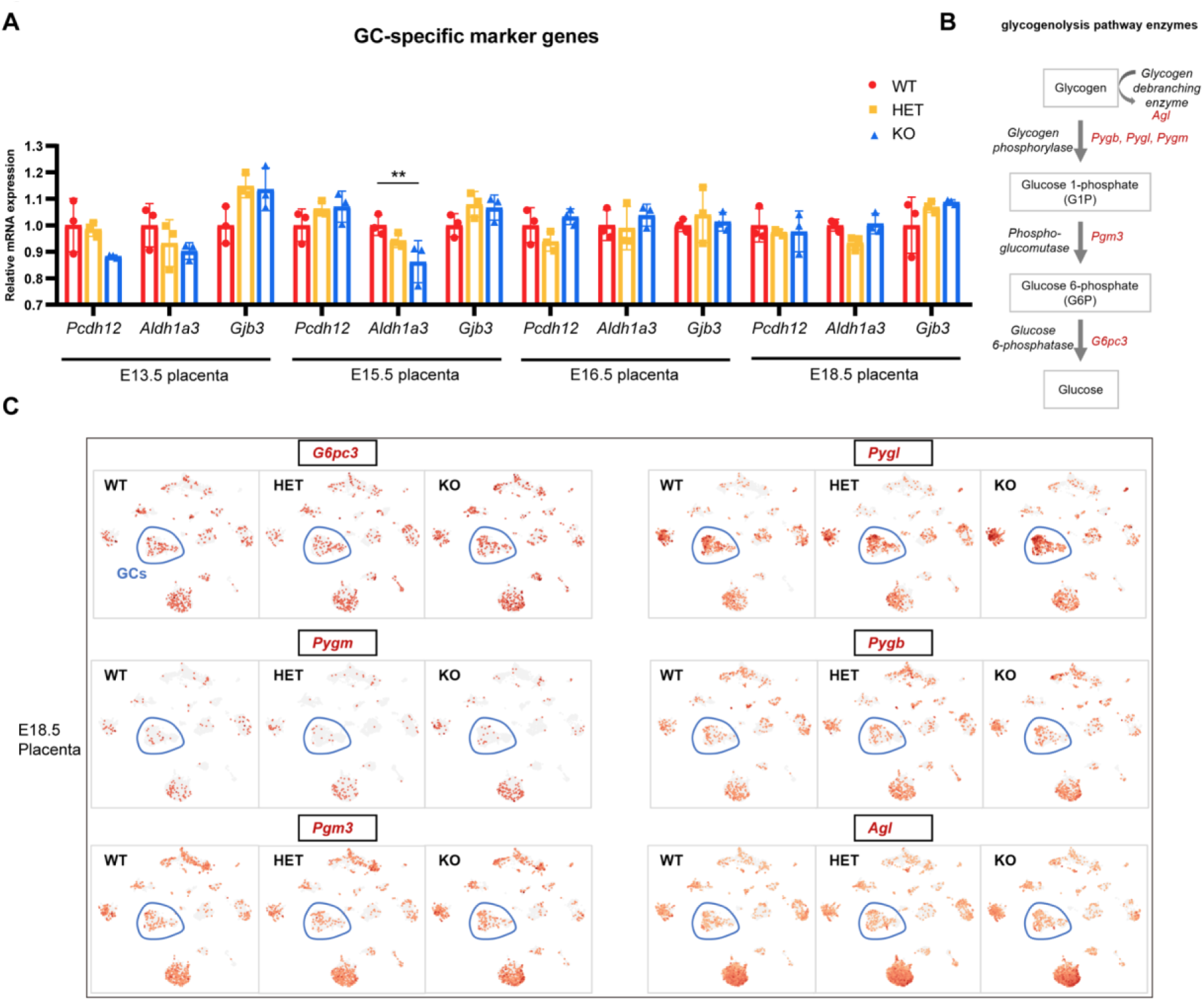
The expression of GC markers and genes encoding key glycogenolytic enzymes by snRNA-seq of E18.5 placentas, related to. **Figure 5** (A) Relative expression of GC-specific marker genes (*Pcdh12*, *Aldh1a3*, and *Gjb3*) in WT, HET, and KO placentas across development. One-way analysis of variance (ANOVA). \*\**p* < 0.01. Data are presented as mean ± SEM. (B) Schematic of glycogenolysis pathway enzymes and the genes that encode them. (C) The expression of genes encoding key enzymes at E18.5 WT, HET, and KO placentas by snRNA-seq data. The blue circles indicate the GC population.

**Figure S10.**
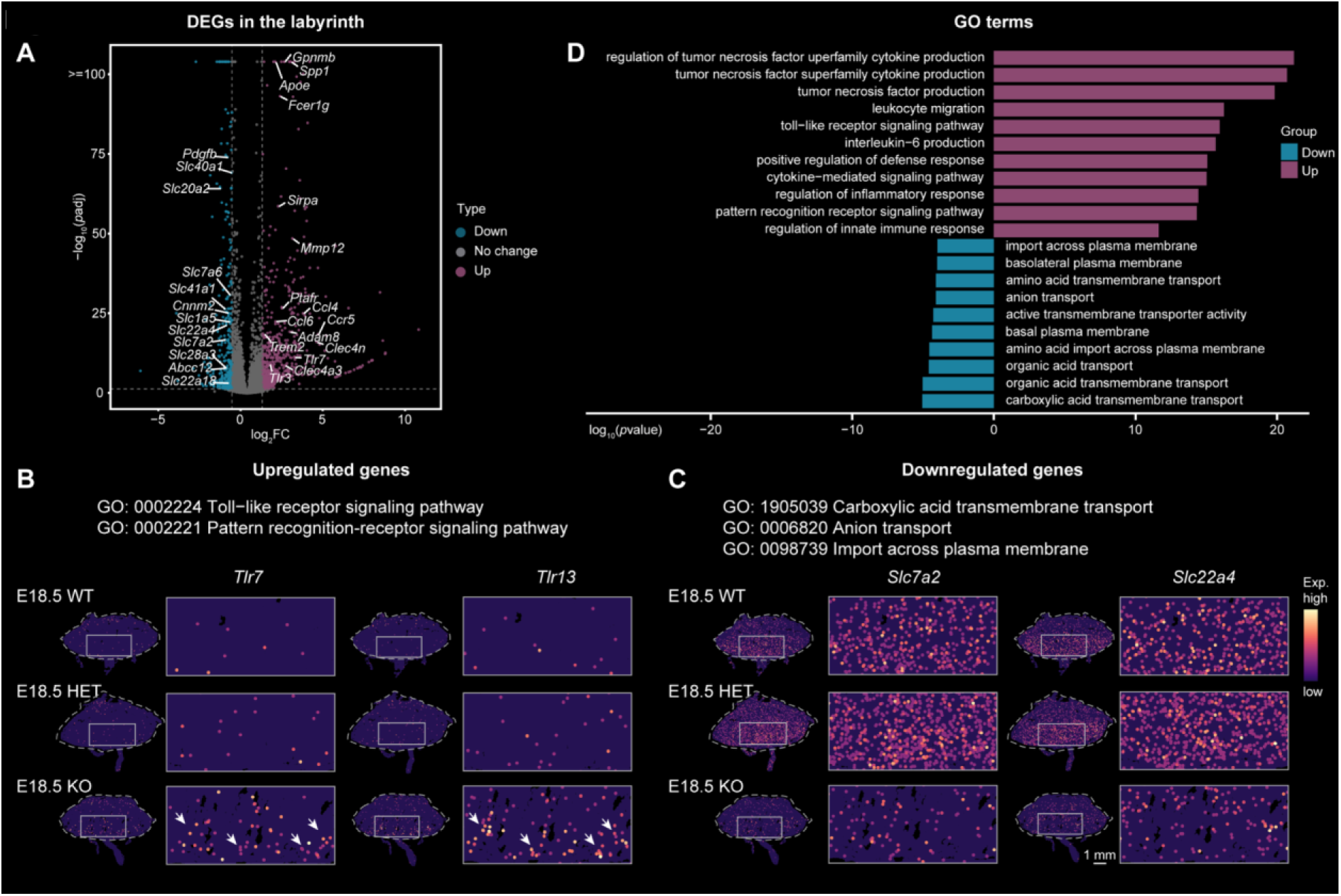
Spatial distribution of differentially expressed genes in the labyrinth between KO and WT placentas, related to. **Figure 6** (A) Volcano plot showing the differentially expressed genes (DEGs) in the labyrinth of KO placentas compared with WT placentas. (B) Spatial visualization of representative upregulated genes in enriched GO terms on the WT, HET and KO placental sections. Scale bars, 1 mm. (C) Spatial visualization of representative downregulated genes in enriched GO terms in the WT, HET and KO placenta sections. Scale bars, 1 mm. (D) Two-sided bar plot showing the enriched GO terms of differentially upregulated and downregulated genes in the labyrinth of KO placentas.

**Figure S11.**
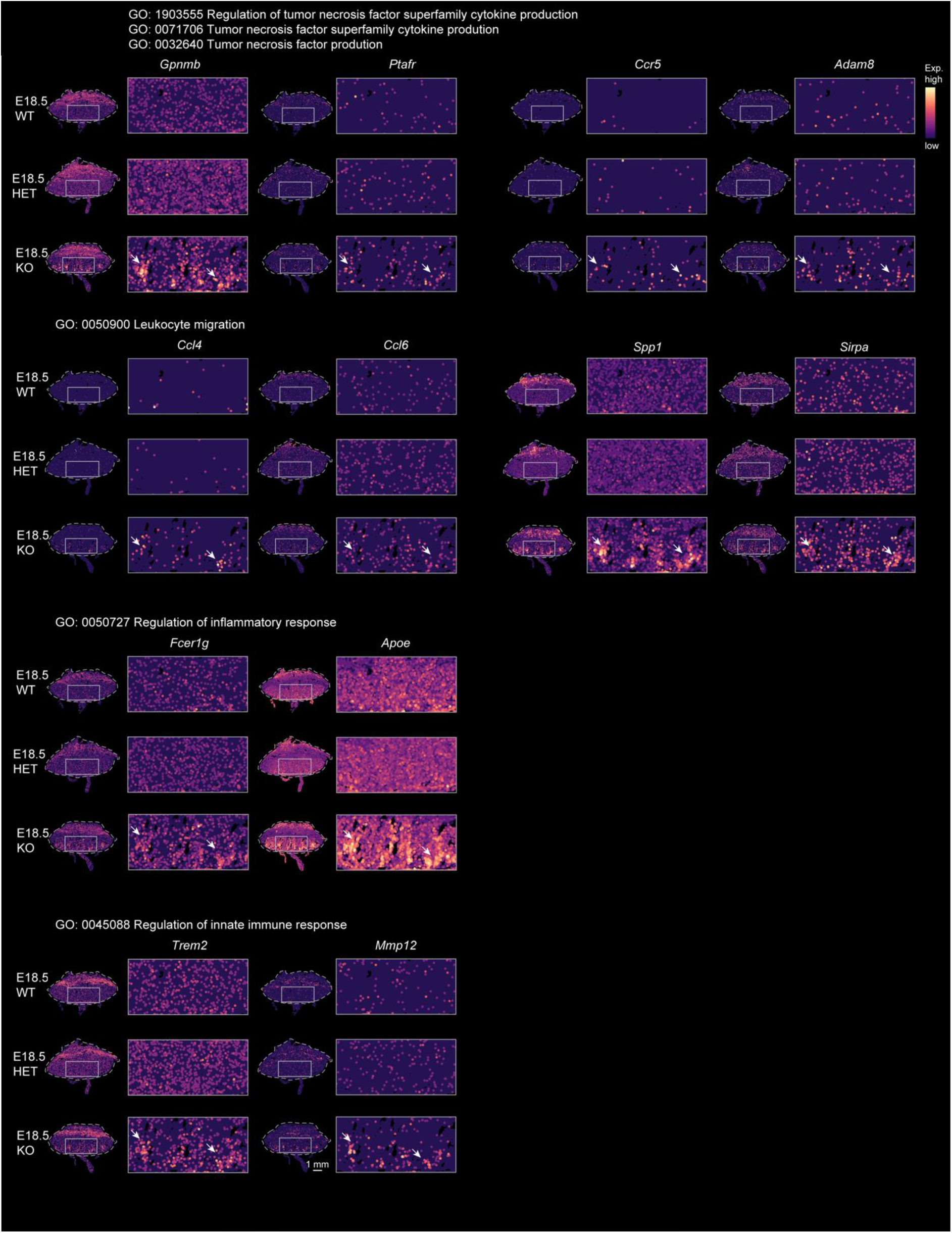
Spatial visualization of unregulated genes in the labyrinth of KO placentas compared with WT placentas, related to. **Figure 6** Spatial visualization of representative upregulated genes in enriched GO terms on the WT, HET and KO placental sections. Scale bars, 1 mm.

**Figure S12.**
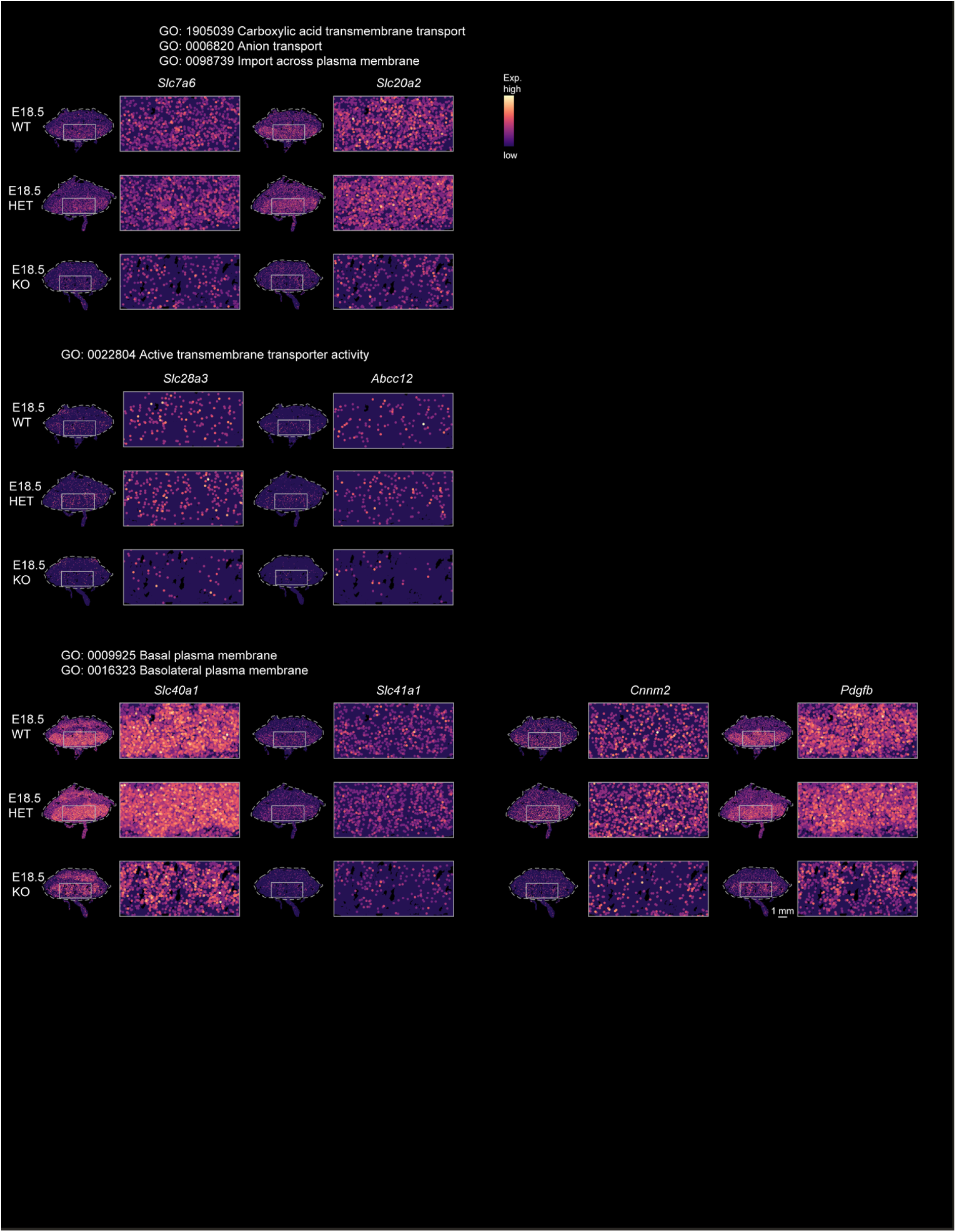
Spatial visualization of downregulated genes in the labyrinth of KO placentas compared with WT placentas, related to. **Figure 6** Spatial visualization of representative downregulated genes in enriched GO terms on the WT, HET and KO placental sections. Scale bars, 1 mm.

**Figure S13.**
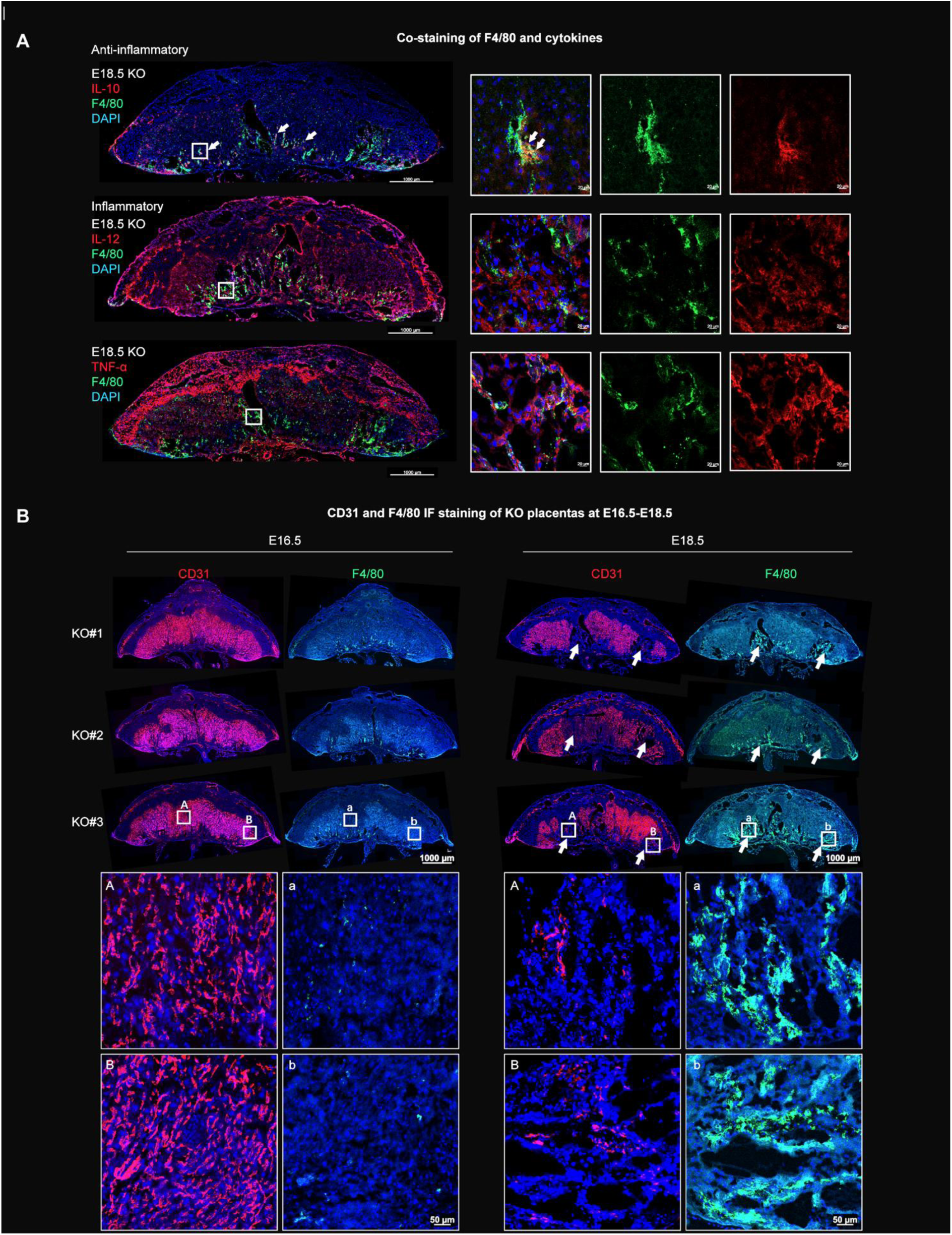
Aberrantly localized macrophages are closely linked to the emergence of vascular abnormalities, related to. **Figure 6** (A) Co-immunostaining of F4/80 and anti-inflammatory cytokine (IL-10), inflammatory cytokines (IL-12 and TNF-α). Scale bars, 1000 μm (overall shape) and 20 μm (high-magnification view). (B) Immunostaining of CD31 and F4/80 on adjacent placental sections. Scale bars, 1000 μm (overall shape) and 50 μm (high-magnification view).

### Supplementary Tables

**Supplementary Table 1. DEGs in all cell types, related to Fig. 1**.

All differential expressed genes (DEG) of all placental and maternal cells.

**Supplementary Table 2. DEGs in trophoblast cells, related to Fig. 2**.

Sub-clustering of trophoblast cells, and all differential expressed genes (DEG) of all trophoblast subtypes.

**Supplementary Table 3. DEGs in GC subclusters, related to Fig. 2**.

All differential expressed genes (DEG) between two GC subtypes, GC-1 and GC-2. **Supplementary Table 4. GO enrichment analysis of GC subclusters, related to Fig. 2**. GO enrichment of DEGs in two GC subtypes, GC-1 and GC-2.

**Supplementary Table 5. DEGs in the labyrinth of KO placenta and GO enrichment analysis, related to Fig. 5**.

All differential expressed genes (DEG) and their enriched GO terms in the labyrinth of KO placenta compared to WT placenta (adjusted *p* < 0.05, |log₂FC| > 1).

**Supplementary Table 6. DEGs in the labyrinthine macrophages of KO placenta and GO analysis, related to Fig. 5**.

All differential expressed genes (DEG) and their enriched GO terms in the labyrinthine macrophages of KO placenta compared to WT placenta (adjusted p < 0.05, |log₂FC| > 1).

**Supplementary Table 7. Primers and immunostaining antibodies.**

Information regarding the PCR and qRT-PCR primers and antibodies used for immunostaining in this study.

