## Supplementary material for "A spatiotemporal transcriptomic atlas of the mouse placenta reveals glycogen cell-mediated metabolic support essential for fetal viability": Tables (Table1-3)

Table 1. All lethal mouse knockout lines according to previous reports.

| | Paper | Gene | Viability P14 | | | Embryonic viability | | | | | | Placenta |
|---|---|---|---|---|---|---|---|---|---|---|---|---|
| | | | Lethal | Subviable | Viable | Before E8.5 Lethal | E9.5-E14.5 Lethal | E14.5 Subviable | E14.5 Viable | Perinatal | Not certain | |
| 1 | Zhang et al., 2020 | Ano6 | | ✓ | | | | | | ✓ | | Yes |
| 2 | Tetzlaff et al., 2004 | Fbxw7 | ✓ | | | | ✓ | | | | | Yes |
| 3 | Araki et al., 2015 | Cul3 | ✓ | | | | | | | | ✓ | N/A |
| 4 | Lu et al., 1997 | Pkd1 | ✓ | | | | | | ✓ | | | Yes |
| 5 | O'Neill et al., 2011 | Prkab1 | ✓ | | | | | | | | ✓ | N/A |
| 6 | O'Neill et al., 2011 | Prkab2 | ✓ | | | | | | | | ✓ | N/A |
| 7 | YANT et al., 2003 | Gpx4 | ✓ | | | ✓ | | | | | | N/A |
| 8 | Ozaki et al., 2015 | Kdm4c | ✓ | | | | | | | | ✓ | N/A |
| 9 | Roth et al., 2012 | Krt1 | ✓ | | | | | | ✓ | | | N/A |
| 10 | Ji et al., 2017 | Smarcd2 | ✓ | | | | | | | | ✓ | N/A |
| 11 | Meier et al., 2018 | Smn1 | ✓ | | | | ✓ | | | | | N/A |
| 12 | Adissu et al., 2014 | Adam17 | | ✓ | | | | | ✓ | | | N/A |
| 13 | Ho et al., 2017 | Apela | ✓ | | | | ✓ | | | | | Yes |
| 14 | Daniel et al., 2012 | Atm | ✓ | | | | ✓ | | | | | N/A |
| 15 | Hoggatt et al., 2013 | Foxf1 | ✓ | | | | | ✓ | | | | N/A |
| 16 | Yuan et al., 2013 | Gcm2 | | ✓ | | | | | | ✓ | | N/A |
| 17 | Merkle et al., 2012 | Gpi | | | ✓ | | | | | ✓ | | N/A |
| 18 | Pyrgaki et al., 2011 | Grhl2 | ✓ | | | | ✓ | | | | | N/A |
| 19 | Bush et al., 2013 | H3f3b | | | ✓ | | | | ✓ | | | N/A |
| 20 | Rix et al., 2011 | Ift80 | ✓ | | | | ✓ | | | | | N/A |
| 21 | Zhu et al., 2018 | Lman1 | | ✓ | | | | | ✓ | | | N/A |
| 22 | Maue et al., 2012 | Npc1 | | ✓ | | | | | | ✓ | | N/A |
| 23 | Hasegawa et al., 2012 | Pip5k1a | | | ✓ | | | | ✓ | | | N/A |
| 24 | Amable et al., 2011 | Ppp5c | ✓ | | | | | | ✓ | | | N/A |
| 25 | Whitlock et al., 2009 | Rock2 | | ✓ | | | | | | ✓ | | N/A |
| 26 | Drapeau et al., 2014 | Shank3 | | | ✓ | | | | ✓ | | | N/A |
| 27 | Katsuragi et al., 2013 | Bcl11b | ✓ | | | | | | | ✓ | | N/A |
| 28 | Langford et al., 2018 | Ly6e | ✓ | | | | ✓ | | | | | Yes |
| 29 | Shu et al., 2002 | Hoxa2 | | | ✓ | | | | ✓ | | | N/A |
| 30 | Morin et al., 2006 | Yap1 | ✓ | | | ✓ | | | | | | N/A |
| 31 | Zhou et al., 2012 | Mns1 | | ✓ | | | | | | | ✓ | N/A |
| 32 | Senkevitch et al., 2012 | Nags | ✓ | | | | | | | ✓ | | N/A |
| 33 | Bainbridge et al., 2011 | Gcm1 | ✓ | | | | ✓ | | | | | Yes |
| 34 | Mittag et al., 2007 | Pax8 | | ✓ | | | | | ✓ | | | N/A |
| 35 | Planells et al., 2000 | Scn2a | ✓ | | | | | | | ✓ | | N/A |
| 36 | Marians et al., 2002 | Tshr | ✓ | | | | | | | ✓ | | N/A |
| 37 | Chen et al., 2012 | Tmeff2 | | | ✓ | | | | ✓ | | | N/A |
| 38 | Zhong et al., 1999 | Tarbp2 | | ✓ | | | | | | ✓ | | N/A |
| 39 | ISHII et al., 2001 | Stambp | | | ✓ | | | | ✓ | | | N/A |
| 40 | Zhou et al., 2000 | Mmp14 | | | ✓ | | | | ✓ | | | N/A |
| 41 | Davisson et al., 2011 | Cntn1 | | ✓ | | | | | ✓ | | | N/A |
| 42 | Laurin et al., 2008 | Dock1 | ✓ | | | | | | | ✓ | | N/A |
| 43 | Yu et al., 2014 | Dph1 | | ✓ | | | | | | ✓ | | N/A |
| 44 | Kojic et al., 2021 | Elp2 | ✓ | | | | ✓ | | | | | N/A |
| 45 | Mostoslavsky et al., 2006 | Sirt6 | | | ✓ | | | | ✓ | | | N/A |
| 46 | Shull et al., 1992 | Tgfb1 | | | ✓ | | | | ✓ | | | N/A |
| 47 | Kruger et al., 2007 | Sp1 | ✓ | | | | ✓ | | | | | N/A |
| 48 | Geiser et al., 2012 | Slc39a4 | ✓ | | | | ✓ | | | | | N/A |
| 49 | Perez et al., 2018 | 9130011E15Rik | ✓ | | | | ✓ | | | | | Yes |
| 50 | | Crb2 | ✓ | | | | ✓ | | | | | Yes |
| 51 | | Ddx42 | ✓ | | | | ✓ | | | | | Yes |
| 52 | | Denr | ✓ | | | | ✓ | | | | | Yes |
| 53 | | Dhodh | ✓ | | | | ✓ | | | | | Yes |
| 54 | | Dpm1 | ✓ | | | | ✓ | | | | | Yes |
| 55 | | Ndufb8 | ✓ | | | | ✓ | | | | | Yes |
| 56 | | Nhlrc2 | ✓ | | | | ✓ | | | | | Yes |
| 57 | | Nubpl | ✓ | | | | ✓ | | | | | Yes |
| 58 | | Pgap2 | ✓ | | | | ✓ | | | | | Yes |
| 59 | | Pigl | ✓ | | | | ✓ | | | | | Yes |
| 60 | | Pitrm1 | ✓ | | | | ✓ | | | | | Yes |
| 61 | | Sqle | ✓ | | | | ✓ | | | | | Yes |
| 62 | | Trub2 | ✓ | | | | ✓ | | | | | Yes |
| 63 | | Vps33b | ✓ | | | | ✓ | | | | | Yes |
| 64 | | Wrap53 | ✓ | | | | ✓ | | | | | Yes |
| 65 | | 1110037F02Rik | ✓ | | | | ✓ | | | | | Yes |
| 66 | | Bap1 | ✓ | | | | ✓ | | | | | Yes |
| 67 | | L3mbtl2 | ✓ | | | | ✓ | | | | | Yes |
| 68 | | Smg1 | ✓ | | | | ✓ | | | | | Yes |
| 69 | | Arhgef7 | ✓ | | | | ✓ | | | | | Yes |
| 70 | | Cnot4 | ✓ | | | | ✓ | | | | | Yes |
| 71 | | Commd10 | ✓ | | | | ✓ | | | | | Yes |
| 72 | | Coq4 | ✓ | | | | ✓ | | | | | Yes |
| 73 | | Crls1 | ✓ | | | | ✓ | | | | | Yes |
| 74 | | Dennd4c | ✓ | | | | ✓ | | | | | Yes |
| 75 | | Dnajc8 | ✓ | | | | ✓ | | | | | Yes |
| 76 | | Eif3h | ✓ | | | | ✓ | | | | | Yes |
| 77 | | Fam21 | ✓ | | | | ✓ | | | | | Yes |
| 78 | | Gpatch1 | ✓ | | | | ✓ | | | | | Yes |
| 79 | | Med23 | ✓ | | | | ✓ | | | | | Yes |
| 80 | | Nadk2 | ✓ | | | | ✓ | | | | | Yes |
| 81 | | Nek9 | ✓ | | | | ✓ | | | | | Yes |
| 82 | | Nrbp1 | ✓ | | | | ✓ | | | | | Yes |
| 83 | | Pigf | ✓ | | | | ✓ | | | | | Yes |
| 84 | | Rab21 | ✓ | | | | ✓ | | | | | Yes |
| 85 | | Setd5 | ✓ | | | | ✓ | | | | | Yes |
| 86 | | Supt3 | ✓ | | | | ✓ | | | | | Yes |
| 87 | | Timmdc1 | ✓ | | | | ✓ | | | | | Yes |
| 88 | | Traf2 | ✓ | | | | ✓ | | | | | Yes |
| 89 | | Kif3b | ✓ | | | | ✓ | | | | | No |
| 90 | | Chtop | ✓ | | | | | ✓ | | | | Yes |
| 91 | | Dhx35 | ✓ | | | | | ✓ | | | | Yes |
| 92 | | Fam160a1 | ✓ | | | | | ✓ | | | | Yes |
| 93 | | Fryl | ✓ | | | | | ✓ | | | | Yes |
| 94 | | Anks6 | ✓ | | | | | ✓ | | | | No |
| 95 | | Ift140 | ✓ | | | | | ✓ | | | | No |
| 96 | | Otud7b | ✓ | | | | | ✓ | | | | No |
| 97 | | Atp11a | ✓ | | | | | | ✓ | | | Yes |
| 98 | | Brd2 | ✓ | | | | | | ✓ | | | Yes |
| 99 | | Cir1 | ✓ | | | | | | ✓ | | | Yes |
| 100 | | Exoc3l2 | ✓ | | | | | | ✓ | | | Yes |
| 101 | | H13 | ✓ | | | | | | ✓ | | | Yes |
| 102 | | Kif1bp | ✓ | | | | | | ✓ | | | Yes |
| 103 | | Psph | ✓ | | | | | | ✓ | | | Yes |
| 104 | | Pth1r | ✓ | | | | | | ✓ | | | Yes |
| 105 | | Rpgrip1l | ✓ | | | | | | ✓ | | | Yes |
| 106 | | Slc25a20 | ✓ | | | | | | ✓ | | | Yes |
| 107 | | Smg9 | ✓ | | | | | | ✓ | | | Yes |
| 108 | | Ssr2 | ✓ | | | | | | ✓ | | | Yes |
| 109 | | 1700067K01Rik | ✓ | | | | | | ✓ | | | No |
| 110 | | 4933434E20Rik | ✓ | | | | | | ✓ | | | No |
| 111 | | Adamts3 | ✓ | | | | | | ✓ | | | No |
| 112 | | Atg16l1 | ✓ | | | | | | ✓ | | | No |
| 113 | | Celf4 | ✓ | | | | | | ✓ | | | No |
| 114 | | Chst11 | ✓ | | | | | | ✓ | | | No |
| 115 | | Cmip | ✓ | | | | | | ✓ | | | No |
| 116 | | Crim1 | ✓ | | | | | | ✓ | | | No |
| 117 | | Cyfip2 | ✓ | | | | | | ✓ | | | No |
| 118 | | Cyp11a1 | ✓ | | | | | | ✓ | | | No |
| 119 | | D930028M14Rik | ✓ | | | | | | ✓ | | | No |
| 120 | | Dmxl2 | ✓ | | | | | | ✓ | | | No |
| 121 | | Fam46c | ✓ | | | | | | ✓ | | | No |
| 122 | | Fut8 | ✓ | | | | | | ✓ | | | No |
| 123 | | Nxn | ✓ | | | | | | ✓ | | | No |
| 124 | | Polb | ✓ | | | | | | ✓ | | | No |

| | | | | | | | |
|---|---|---|---|---|---|---|---|
| 125 | Prrc2b | ✓ | | | | ✓ | No |
| 126 | Sh3pxd2a | ✓ | | | | ✓ | No |
| 127 | Slc5a7 | ✓ | | | | ✓ | No |
| 128 | Tcf7l2 | ✓ | | | | ✓ | No |
| 129 | Traf6 | ✓ | | | | ✓ | No |
| 130 | Trim45 | ✓ | | | | ✓ | No |
| 131 | Gtpbp3 | | ✓ | ✓ | | | Yes |
| 132 | Camsap3 | | ✓ | | ✓ | | Yes |
| 133 | Prmt7 | | ✓ | | ✓ | | No |
| 134 | Adcy9 | | ✓ | | | ✓ | Yes |
| 135 | Cfap53 | | ✓ | | | ✓ | Yes |
| 136 | Col4a3bp | | ✓ | | | ✓ | Yes |
| 137 | Actn4 | | ✓ | | | ✓ | No |
| 138 | Arid1b | | ✓ | | | ✓ | No |
| 139 | Capza2 | | ✓ | | | ✓ | No |
| 140 | Cpt2 | | ✓ | | | ✓ | No |
| 141 | Dbn1 | | ✓ | | | ✓ | No |
| 142 | Gm5544 | | ✓ | | | ✓ | No |
| 143 | Hmgxb3 | | ✓ | | | ✓ | No |
| 144 | Mybphl | | ✓ | | | ✓ | No |
| 145 | Nsun2 | | ✓ | | | ✓ | No |
| 146 | Pdzk1 | | ✓ | | | ✓ | No |
| 147 | Rala | | ✓ | | | ✓ | No |
| 148 | Rundc1 | | ✓ | | | ✓ | No |
| 149 | Smpd4 | | ✓ | | | ✓ | No |
| 150 | Syt1 | | ✓ | | | ✓ | No |
| 151 | Unk | | ✓ | | | ✓ | No |

Table 2. Genotype distribution of Ano6 knockout mice.

| | Total | WT | HET | KO |
|---|---|---|---|---|
| **Ano6 deletion** | 132 | 40 (30.3%) | 88(66.67%) | 4 (3.03%) |

Table 3. Genotype distribution of Ano6 knockout mice after glucose supplementation with pregnant mice.

| | Total | WT | HET | KO |
|---|---|---|---|---|
| **Glucose supplement** | 74 | 19 (25.7%) | 47 (63.5%) | 8 (10.8%) |
